# Computational Stabilization of the Human VH Germline Repertoire to Enable Conditional Multi-Specific Therapeutic Development

**DOI:** 10.1101/2025.10.31.685826

**Authors:** David F Thieker, Jayd Hanna, Pricilla White, Glenn Capodagli, Christen Buetz-Duncan, Christina D Carnevale, Arlene Sereno, Falene Chai, Abby Lin, Craig R Pigott, Rajesh K Sharma, Natasha Del Cid, Nathan I Nicely, Brian Kuhlman, Stephen J Demarest

## Abstract

Protein-based biologic therapies, particularly antibody-like therapeutics, have emerged as a major modality to treat nearly all chronic and infectious diseases. Antagonists have dominated the first wave of antibody and antibody-like biologics, whereas agonism has generally been challenging due to systemic activation leading to issues with therapeutic index and problems with pleiotropic activity. Additionally, agonists that use native proteins such as cytokines can be challenging to produce at scale given these proteins evolved to act locally and are not designed for large scale manufacturing. To address these challenges, we generated a biologics platform comprised of stabilized human VH domains (VH-Select™) which encompass the entire human germline repertoire with the goal of building multispecific biologics denoted Tentacles™ that use avidity-based binding to achieve conditional activity directed to specific cell types or tissues. Stabilizing disulfides and point mutations were identified computationally with Rosetta, evaluated *in vitro*, and combined into designs with 3-5 amino acid substitutions for each of the seven germline families (VH1-VH7). Computational design was also employed to reduce dimerization from both VL and homotypic VH-VH interactions. Optimization of specific sequences improved expression by greater than 600-fold and thermostability by more than 20°C. Each of the germline variants were screened for low HLA class II binding and incorporated into a library which demonstrated significant improvements in cellular protein production, thereby increasing sequence diversity for screening campaigns. These VH-Select™ scaffolds are useful for the discovery of novel binders used to build multispecific Tentacles™ designed for cis-interactions that achieve cell- and tissue-specific activation. As an example of the utility of the platform, we generated a set of Tentacles™ that use VH-Select™ binders to conditionally agonize IL2Rγβ and 41BB on PD1+, LAG3+, or CD8+ T cells. These Tentacles™ demonstrate promising manufacturability, antibody-like exposure *in vivo*, and strong anti-tumor activity in a humanized tumor model. Overall, we believe the incorporation of VH-Select™ binders into multispecific Tentacles™ has the potential to create a host of conditionally active biologics to treat various chronic and acute diseases.

## Introduction

Based on the vast diversity generated by VDJ recombination, antibody VH domains drive the primary specificity of most antibodies ^1^. Given VH domains are ten-fold smaller than IgGs, their use as binding scaffolds provides many advantages ^2,3^. Their small size allows for increased tissue penetrance ^4,5^. An evolved or engineered ability to bind without light chains also conveys modularity that is difficult to achieve with natural IgGs due to obstacles of multi-chain assembly ^6^. Along with repertoires of normal antibody VHs and VLs, camelids evolved ‘Variable Heavy domains of the Heavy chain’ (VHH). These sequences contain VH-like moieties with numerous framework and CDR modifications which enable intrinsic stability and solubility improvements, along with the elimination of VL interactions ^7^. The use of VHH domains as biotherapeutics generally requires a balance between humanization and maintenance of key non-human germline residues, which can pose an immunogenicity risk for their use. The ideal solution is the use of fully human VH domains for discovery ^8^.

Human VH domains, however, evolved in the context of the antibody Fab (Fragment of antigen binding), which comprises the heterodimeric complex of VH-CH1 from a heavy chain (HC) and VL-CL domains from a light chain (LC). Isolating VH domains from this complex commonly results in proteins with poor biophysical characteristics. First, isolated VHs have midpoints of thermal unfolding (Tm) approximately 20-25 °C lower than what is observed in the Fab context ^9–12^. Unlike camelid VHHs, human VH domains tend to irreversibly aggregate upon unfolding in normal aqueous buffers ^13,14^. When expressed in eukaryotic cells at 37 °C, the low Tm of VHs leads to a significant population of intrinsically unfolded material and a continual and irreversible march towards proteolysis and/or aggregation. An additional liability of the human VH scaffold is that it has a propensity to bind the VL domain from free light chains as well as form homodimers. VH/VL and VH/VH dimerization propensity can be strong or weak depending on the germline frameworks and CDRs with lambda VLs demonstrating a higher affinity towards human VHs than kappa VLs ^9,12^. High resolution structures of two human VH domains independently isolated using phage display were found to be dimers ^15,16^. Interestingly, dimerization in these structures used VH residues at the canonical VL interface, but with different side chain orientations that were nearly identical at the VH-VH interface within the two structures.

Significant efforts have been made to isolate or engineer VH domains with improved biophysical properties ^17^. These efforts have included consensus engineering and transfer of optimal framework residues to alternate frameworks ^12^, phage display using conformational protein A binding or heat treatments to pan for improved biophysical properties ^14,18–20^, identification of single ideal VH domains for discovery ^3^, or the addition of stabilizing disulfides ^11,21–24^. Stabilizing modifications often use residues within the CDRs of single VH frameworks for the creation of *in vitro* screening libraries. Screening using a single VH framework has limitations in the paratopes available for recognizing different epitope topologies, which consist of diverse polar, hydrophobic, charged surface compositions. Modifying HCDR1 and HCDR2 to add non-native amino acids that expand the paratope space of a single VH or a small family of VHs runs the risk of introducing MHC II epitopes and immunogenic potential.

Here, we strove to generate non-immunogenic stabilizing VH designs that would enable use of the intrinsic paratope diversity afforded by the entire human VH germline repertoire. We used a computational approach to identify germline-specific stabilizing designs without incorporating strong MHC II epitopes. Ultimately, every functional human VH germline in the human genome was stabilized by making 3-5 modifications per germline family that resulted in antibody-like expression and stability. Additionally, mutations were identified that reduced intrinsic VH/VL and VH/VH dimerization, enabling improved modularity without interdomain association. We denote these stabilized VH germlines as VH-Select™. We then demonstrate the ability of these VH-Select™ modifications to improve the expression properties of diverse VH libraries; show their utility for generating conditionally active multispecifics (Tentacles™) with antibody-like stability and *in vivo* exposure; and report a Tentacle™ that enabled strong anti-tumor activity without inducing the systemic toxicity observed for the untargeted agonists.

## Materials and Methods

### Computational disulfide bond design

Homology models were created for six diverse VH sequences selected from previous HuTARG wild-type VH discovery campaigns by identifying the most suitable crystal structures (considering resolution and sequence similarity) and mutating the residues to match with Rosetta’s side chain packing protocol ^25^ (**{Table S1**). The VH coordinates were all originally complexed within the multidomain context of an antibody Fab. Atoms within the VH domains were extracted and used as templates for homology models. The starting VH structures were diversified by building two homology models from separate crystal structures for most of the selected sequences (**Table S1**). The energies of these homology models were then minimized within the Rosetta software using a combination of AtomTree and Cartesian minimizations ^25,26^. Computational prediction of stabilizing disulfide bonds was performed by cycling through all residue pairs, mutating both residues in the pair to Cys, and evaluating the favorability of a potential disulfide bond with the Rosetta energy function^27^. The results were sorted based on the disulfide score (dslf_fa13) and models scoring less than −0.3 were considered for experimental testing.

### Computational saturation mutagenesis

Computational design was also applied to predict stabilizing point mutations. The same energy-minimized homology models used for disulfide prediction were used to create libraries of predominantly single variants, along with a small number of combinatorial variants, that may improve stability. Site saturation mutagenesis, in which each position within the protein was replaced with all possible amino acids (excluding Cys), was performed with previously described RosettaScripts protocols ^28,29^. The energy of each point mutation was compared to the score of the WT sequence to calculate the difference in energy (ΔE). The average score for both homology models of the target sequence were then sorted by value to rank the mutations. *In silico* immunogenicity testing was performed using the NetMHCIIpan 4.3 EL algorithm available at the Immune Epitope Database and Tools (IEDB) website (https://iedb.org/) ^30^. Peptides were evaluated against the 7 most common class II HLA alleles and were constructed with each single mutation at the center of a 17-residue epitope (or double mutation if the two were within 9 amino acids of one another) ^31,32^. Scores <10 are considered weakly immunogenic and scores <1 are considered highly immunogenic. No scores below 1 were found and scores between 1 and 10 were typically limited to a single HLA II allele. Results are provided in Supplementary File NetMHCIIpan 4.3 IEDB Scoring.xlsx.

Assessment of the impact of mutations at the VH/VL and VH/VH interface were also performed using ThermoMPNN ^33^. To uncover mutations that significantly impair VH/VL and VH/VH interactions without impacting the stability of the VH monomer, ThermoMPNN stability assessments of 19 amino acid substitutions at every position of the VH domain were performed on the heterodimers, homodimers, and monomers (VH/VL = 4LLU, 7VYT; VH/VH = 3QYC, 1OHQ).

### Crystallography

The anti-IL2Rγc VH3-20 protein was expressed with an octa-histidine tag and purified via standard Ni^2+^-NTA procedures then further purified via size exclusion chromatography using a Superdex 200 prep grade 16/60 (Cytiva) column with a running buffer of 20 mM HEPES pH 7.4, 150 mM NaCl. Peak fractions were collected and the pool concentrated to 11.9 mg/ml as determined by A280.

The anti-IL2Rγc VH3-20 protein was tested for crystallization against common commercially available crystal screens using a Mosquito dropsetter (SPT Labtech) with drops composed of 200 nl protein and 200 nl reservoir solution set over 30 ul reservoir volumes. Crystals were observed in eight days over a reservoir solution composed of 0.2 M calcium acetate, 0.1 M MES pH 6.0, 20% PEG 8000. The crystals were briefly soaked in reservoir supplemented with 15% ethylene glycol then cryocooled in liquid nitrogen. Diffraction data were collected at Southeast Regional Collaborative Access Team (SER-CAT) 22-ID beamline at the Advanced Photon Source, Argonne National Laboratory, using an incident beam of 1 Å in wavelength. Data were reduced in HKL-2000^34^. The structure was phased by molecular replacement using Phaser^35^ with PDB 7JOO as the search model^36^. Real space rebuilding were done in Coot^37^, and reciprocal space refinements and validations were done in PHENIX^38^. Coordinates and structure factors have been deposited in the Protein Data Bank (PDB) with accession number <u>9PNX</u>.

### Computational modeling of stabilizing designs

Models of VH1-69.2, VH2-26, VH4-39, and VH5-51 were generated using AlphaFold2^39^. Protein structure figures were generated using PyMol (https://www.pymol.org). Electrostatic mapping of the surfaces of these proteins was performed in PyMol using the APBS Electrostatics plugin. Amino acid dihedral distributions were assessed using the Top8000 dihedral library database in MolProbity ^40,41^. Generation of Ramachandran plots was performed based on Ramachandran_Plotter ^42^.

### Discovery of IL2Rγ, IL2Rβ, and 41BB VH binders

Discovery of IL2Rγ, IL2Rβ, and 41BB VH binders was performed using the HuTARG^TM^ mammalian display system ^43^. Briefly, the most predominant 47 functional human germline VH domains (−01* allele at IMGT^44^) were included within the HuTARG library as membrane tethered proteins. IL2Rγ (CD132), IL2Rβ (CD122), and CD8a soluble extracellular domains were obtained from R&D Systems. The antigens were biotinylated using standard kits and lysine chemistry. The mammalian display library was expanded and screened for binding to these antigens by flow cytometry using streptavidin-PE as a detection reagent. Typically, two rounds of FACS enrichment followed by expansion were needed to obtain a solid pool of VH binders whose sequences could be identified using next generation sequencing (NGS) on a MiSEQ device (Illumina). Discovery of the active IL2Rγ/IL2Rβ VH pairing was described in a patent filing ^45^.

### Protein Production and Testing

Cloning and plasmid production were performed using standard molecular biology methods. All VH domains were produced with either a human IgG1-Fc or with a C-terminal octa-histidine tag into a plasmid containing a CMV promotor driven open reading frame and BGH polyA tail. Secretion was driven using a mouse IgG signal peptide. Secreted proteins were produced by transfecting plasmids into HEK293 cells for transient expression using the Thermofisher Expi293 system. Supernatants for protein characterization were collected via centrifugation and then filtered. Titer determination was performed on a GatorBio biolayer interferometry (BLI) instrument using anti-Histidine Tag Tips for VH-Histag domains and Protein A Tips for VH or multi-specific proteins containing a human antibody Fc with purified VH-Histag and VH-Fc proteins used as standards, respectively. For stability measurements, mammalian supernatants were analyzed using differential scanning fluorimetry (DSF) using a QuantStudio3 according to the manufacturer’s protocols (Applied Biosystems) and the fluorescence vs temperature curves were analyzed using Applied Biosystem’s Protein Thermal Shift^TM^ software version 1.4. Analytical SEC measurements were performed on a Vanquish Flex UPLC system using a 7.8 x 300 mm Mab-PAC analytical HPLC column (QS065397-EN) and Cell Mosaic protein standards (CM92005).

### Light Chain Pairing Assay with VH Variants

HEK293 cells were seeded at a density of 0.5 × 10^6^ cells per well in 24-well plates (Greiner #662102) in Freestyle F17 media (Gibco #A1383502) + 10% fetal bovine serum (Gibco #12483020) + 1% Penicillin/Streptomycin (Gibco #15070063) + 1% L-Glutamine (Gibco #25030081). These cells were transfected with VH-Fc-transmembrane (PDGFR) variants with mutations to reduce light chain pairing and a common kappa light chain. 1.2 μg of PEI Max Hydrochloride (Polysciences #24765) was diluted in Opti-MEM media (Life Technologies #31985088) with 0.4 μg of plasmid DNA for 20 minutes before adding to cells. Transfected cells were left to incubate for 24 hours at 37°C with 5% CO_2_ on a 150 RPM shaking platform. Binding of kappa light chain to VH-Fc-transmembrane variants was determined by flow cytometry on a BD FACS Celesta using Goat anti-Human Kappa light chain R-PE (Southern Biotech #24765) at 0.1 μg/mL concentration and AlexaFluor 647 Affinipure Goat anti-Human IgG Fcγ Fragment Specific (Jackson ImmunoResearch #109-605-098) at 0.3 μg/mL concentration. A normalized expression gate was applied to all samples and a geomean fluorescence of kappa light chain binding was analyzed.

### Phospho-STAT5 flow cytometry assay

Briefly, PBMCs are isolated from leukopacs collected at the San Diego blood bank and frozen. After thawing, the PBMCs are stimulated for 48 hours in a flask coated with anti-CD3 (OKT3, Biolegend) and with soluble anti-CD28 (CD28.2, Biolegend) to induce PD1 and LAG3 expression, then rested overnight in the absence of stimulation. Activation of the PBMCs by Tentacle™ test articles was performed for 15 minutes at 37 °C. The cells are then incubated with anti-CD3 (SK7, Biolegend), anti-CD8 (SK1, Biolegend), anti-CD4 (SK3, Biolegend), anti-CD56 (5.1H11, Biolegend), and a fixable viability dye (Zombie Aqua, Biolegend) for 20 minutes on ice, washed, and fixed with BD CytoFix Fixation Buffer (BD Sciences). The cells are then permeabilized with BD Phosflow Perm Buffer III (BD Biosciences), washed, and stained for pSTAT5 (47/Stat5, pY694, BD Biosciences). Samples were acquired on the Novocyte3000 (Agilent Technologies), and analyzed using the NovoExpress software (Agilent Technologies).

### 4-1BB reporter assay

4-1BB/NFκB reporter HEK293 cells (BPS Bio #79289) were seeded at a density of 0.8×10^6^ cells per well in 6-well plates (Corning #353046) in MEM medium (Hyclone #SH30024.01) + 10% fetal bovine serum + 1% non-essential amino acid cell culture supplement (Hyclone #SH30238.01) + 1 nM sodium pyruvate (Hyclone #SH30229.01) + 1% penicillin/streptomycin and incubated overnight at 37 °C. For transfection of LAG3 into the reporter line, lipofectamine 3000 (Invitrogen #L3000008) was diluted into Opti-MEM media (Hyclone #31985-062) in the presence or absence of 5 ug plasmid harboring the full-length LAG3 or PD1 sequence. This lipofectamine/plasmid solution was incubated for 10-15 minutes then added to cells in the 6 well plate for 2-3 days at 37 °C. Transfected cells were detached using Accutase (Innovative Cell Tech #AT104) according to manufacturer’s protocol, spun down, and resuspended at 0.35 × 10^5^ cells per well in 96-well white-bottom plates and incubated overnight at 37 °C. Tentacles™ were then added to the wells and incubated for 5-6 hrs at 37 °C. Next, 0.1 mL One-Step Luciferase reagent (BPS Bio #60690-1) was added at room temperature for 20 min and luminescence was measured on an iD5 reader (Molecular Devices). Cell-free control wells were subtracted from the luminescence reading of all wells.

### Pharmacokinetics in C57Bl/6 mice

Pharmacokinetic studies were approved by an IACUC committee containing both Tentarix and Explora veterinary staff. C57BL/6 mice were sorted into groups of 4 and Tentacles™ were administered via intravenous retro-orbital injection at doses of 0.65 mg/kg. Blood was collected at various time points via submandibular bleeds using K2EDTA collection tubes (BD 365974). Blood was centrifuged and plasma was collected for analysis. The concentration of Tentacles™ was assessed using CD8, PD1, or LAG3 (R&D systems) coated to ELISA plates (Corning 2592) and detected using goat anti-human IgG-Fc (AbCam AB97225). Concentrations were determined by generating standard ELISA curves and testing multiple serum dilutions to ensure there were ELISA readings within the linear range of the assays.

### In vivo tumor growth study

All studies were approved by an IACUC committee containing both Tentarix and Explora veterinary staff. HLA-A2+ A375 cells (ATCC, CRL-1619) were transduced to express CMV antigen by Genecopoeia™. NSG-MHC I/II DKO mice (Jackson Laboratories, Strain 025216) were engrafted with 5 million A375-CMV+ cells in their right flank (subcutaneous) and 10 million HLA-A2+/CMV+ human PBMCs (retro-orbital delivery). Tentacles™ or control molecules were administered intraperitoneally (n = 8 per group) every 7 days beginning on day 4 post tumor and PBMC engraftment. Dosing was initiated 4 days after tumor and PBMC engraftment and repeated weekly. Dosages were as follows: IgG Isotype 2 mg/kg; Combined Urelumab/Pembrolizumab/IL15-IL15R-Fc was 5/5/1 mg/kg; PD1 Tentacle 2.5 mg/kg. Weight and tumor volume were monitored throughout the study period. Significant differences between groups were assessed using a 2-way ANOVA with Tukey multiple comparison test.

### Tissue processing and Flow cytometry analysis of engrafted immune cells in blood and tumors of huPBMC/NSG-DKO mice

On day 25, mice were euthanized and blood was collected with K2EDTA tubes and kept on ice, while tumors were collected and processed following the protocol provided with the Tumor Cell Isolation Kit, human (Miltenyi Biotec, 130-108-339). Tumor and blood cells were then surface stained with the following markers from BioLegend: CD45 (clone HI30), CD3 (clone SK7), CD4 (Clone SK3), CD8 (Clone SK1), CD19 (Clone HIB19), CD56 (Clone HCD56), and PD1 (EH12.2H7). Stained samples were run on Agilent multicolor flow cytometer. Data was analyzed using the NovoExpress analysis software from Agilent Technologies.

## Results

### Computational designs to stabilize each VH germline family

Mutations to stabilize the folded structures of VH sequences across germline families were predicted with Rosetta. A set of diverse VH domains with different V-D-J usage were previously discovered using HuTARG (mammalian display) in binding campaigns against various antigens using libraries containing predominantly native germline V-D-J sequences ^46,47^. All VH-alleles were *01, which was typically the most prevalent allele across the population assessed within the IMGT database (VH1-VH7; https://www.imgt.org). Homology models were built for one or more VH domain sequences selected from these discovery campaigns, accounting for five of the seven human VH germline families (**Table S1**). Mutations predicted *in silico* were cloned individually into 14 VH domains spanning across all seven human VH germline families, each containing a unique heavy chain CDR3 derived from V-D-J recombination.

A set of five disulfide bonds were predicted to form across multiple VH domains with different V-D-J germline usage (**Figure 1b**). All five disulfide bonds increased the expression and/or stability of at least one sequence *in vitro* and some were broadly stabilizing across diverse starting sequences (**Table S2**). The most generally stabilizing disulfide differed across germline families with VH1 / VH7 preferring the 17C_82aC disulfide, VH2 preferring the 19C_81C disulfide, and VH3 / VH4 / VH6 families preferring the 23C_77C disulfide (Kabat numbering ^48^). None of the disulfides were suitable for the VH5 germline sequence. The 35C_50C disulfide is commonly found in rabbit antibodies and has been shown to be stabilizing to various human VHs previously ^24^. It was also found to be stabilizing to the majority of human VH domains here but significantly reduced the binding affinity of the VH domains it was placed into (**Figure 1e, g**). The 17C_82aC and 23C_77C disulfides were shown to have little impact on the binding properties of the VH domains (**Figure 1e,g**).

**Figure 1.**
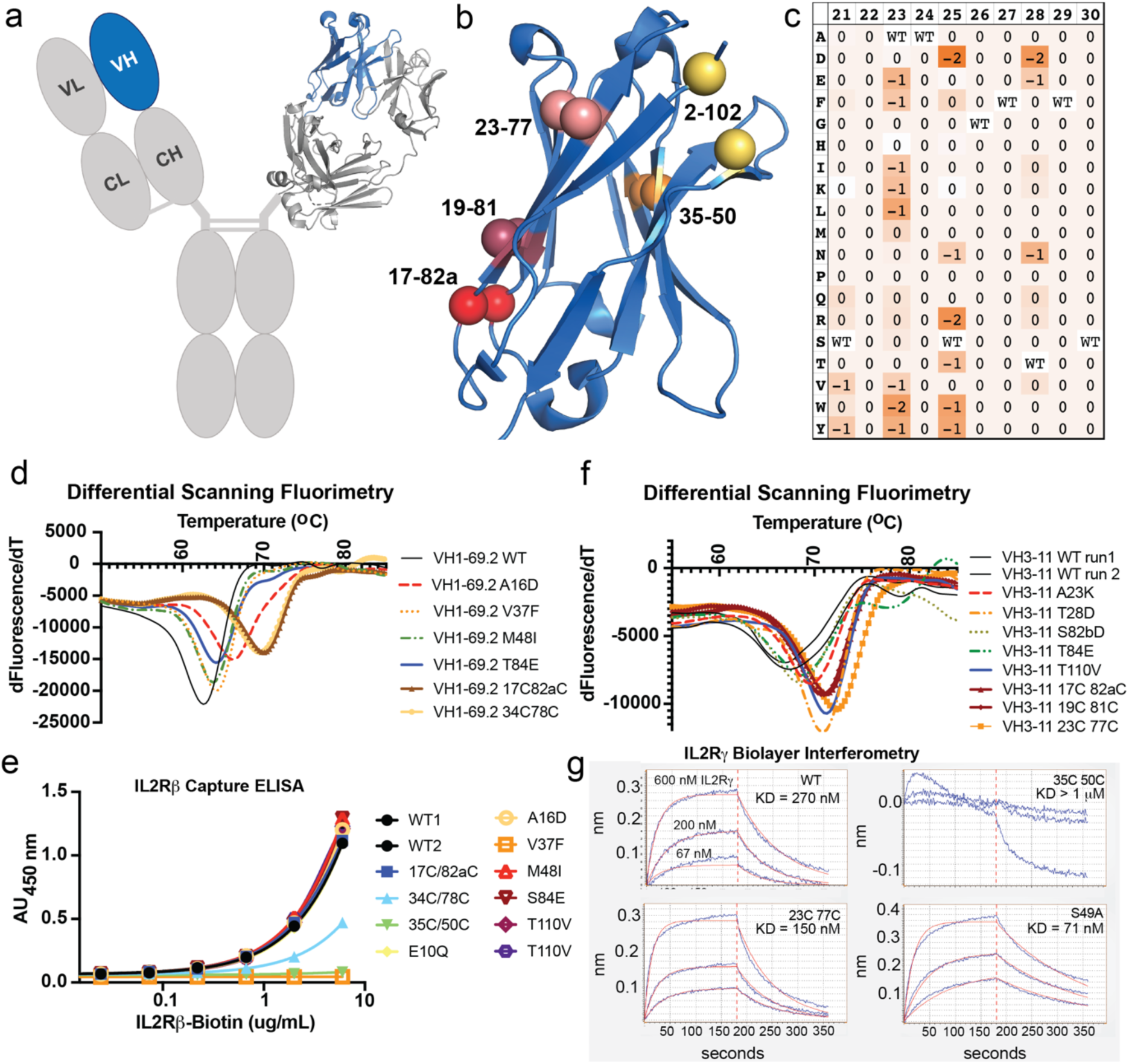
Assessment of disulfide and single mutant impacts on VH stability and binding. Schematic diagram of a VH domain (blue) within an IgG antibody (FAb from PDB: 6J6Y) (a). The extracted VH domain from 6J6Y which depicts the location of the five predicted disulfides that were assessed for their ability to stabilize each germline family. Red, burgundy, salmon, orange, and yellow correspond to disulfides 17C_82aC, 19C_81C, 23C_77C, 35C_50C, and 2C_102C (b). Example of a Rosetta-generated heatmap that depicts the difference in free energy scores between the native sequence and point mutations. Negative values indicate stabilizing amino acid substitutions within the VH domain (c). Representative DSF traces demonstrating the effects of mutation on thermostability (d,f). Relative binding affinity of stabilized VH1 domains recognizing IL2Rβ using ELISA (e) and VH3 domain interactions with IL2Rγ according to biolayer interferometry (BLI; g).

Next, computational design was used to identify point mutations expected to result in a stability increase (**Figure 1c)**. Roughly 30 variants per VH domain were generated, tested for expression, and evaluated for changes to thermal stability via differential scanning fluorimetry (DSF). Variants that increased the expression and/or stability of at least one germline of each VH family (VH1-VH5) are provided in **Table S3**. The VH6-1 and VH7-4 germlines were excluded from the analyses since they only have a single member each. DSF examples of variants increasing the stability of VH1-69.2 and VH3-11 proteins with differing D-J usage are shown in **Figures 1d, f**. The most stabilizing variants for each family were:

VH1: 16D, 48I
VH2: 15G, 44D, 83T
VH3: 49A, 74E, 84E
VH4: 10G, 82bD, 84P
VH5: 28D, 48I, 83D
(The wild-type amino acid is not listed because it may not be conserved across the different germlines of each VH domain).
T110V and T110I were stabilizing across multiple VH families, but in every case led to reduced expression.

While the ultimate goal of the stabilization campaign was to improve the entire set of human germline VH domains as scaffolds for future discovery purposes, and maintaining the existing binding properties of the VH domains was not mission critical, we did evaluate the impact on antigen binding by adding the single amino acid variants as well as the top disulfides to a sequence from the VH1 and VH3 families. Anti-IL2Rβ (VH1-69.2) and anti-IL2Rγ (VH3-20) were discovered using mammalian display with wild-type VH domains ^46^. The VH1.69.2 VH domain binds IL2Rβ with weak affinity precluding the ability to use kinetic measurements; therefore, a capture ELISA was developed to assess binding. The majority of variants bound with similar affinities as the parent molecules; however, the 34C_78C and 35C_50C disulfides, as well as the V37F variant, significantly reduced binding to IL2Rβ (**Figure 1e**). Similar results were obtained for the anti-IL2Rγ VH3-20 variants. Most variants had no impact on IL2Rγ binding; however, addition of the 35C_50C disulfide significantly reduced binding with representative binding curves shown in **Figure 1g**.

### Stabilizing combinations that generally increase the expression and thermal stability of every human VH germline

After identifying disulfides and residue-specific modifications that stabilize the germlines, combinations of these variants were tested for additive or synergistic effects. Immunogenic potential was considered for each 9-mer variant within the combinations using the *in silico* MHC II binding tool at IEDB ^44^ (**Supplementary Data S1**). Peptides that bound to multiple HLA alleles with consensus percentile rankings within the top 1% were culled. Multiple variant combinations were tested in single VH1, VH2, VH4, VH5, VH6, and VH7 proteins and in three separate VH3 family proteins. Variants were selected for combinations based on distal proximities to one another making it likely their stabilizing contributions would be additive. The top designs to emerge that led to increases in protein expression and/or protein stability for each VH protein are provided in **Table 1**. The top design for each family was additionally cloned into a representative VH for every germline family. The impact of the stabilizing designs was assessed by evaluating the expression of each VH domain in comparison to its wild-type counterpart (**Table 2**). Overall, the large majority of germline VHs demonstrated improvements in protein expression compared to their wild-type counterparts (**Figure 2a**). Combination designs also led to significant improvements often between 15-20 °C in the Tm of each VH domain based on DSF measurements (**Figure 2b**). A plot of the impact of the combination designs on the Tms of representative VH(s) from each family (VH1-VH7) is shown in **Figure 2c**. Overall, the majority of VH domains across different germline families demonstrated significant increases in both expression and thermal stability with a small number of mutations (3-4) per domain.

**Figure 2.**
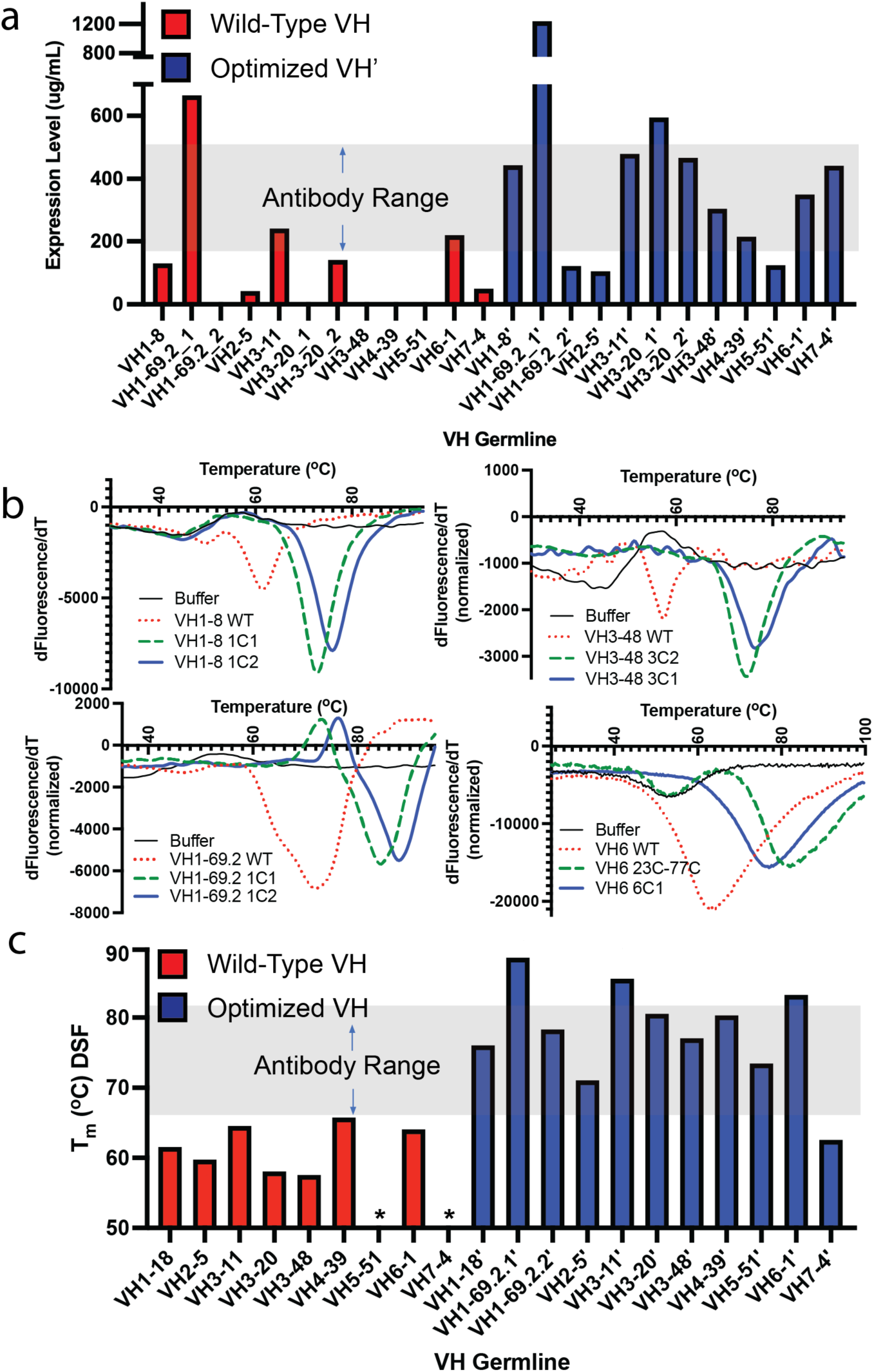
Impact of combining disulfide stabilization with Rosetta based stabilizing mutations. Relative expression levels of non-stabilized (red bars) and stabilized (blue bars) VH domains representing all VH germline families where * indicates a lack of expression (a). DSF of non-stabilized (red dotted) and the top two stabilizing designs (green dotted and blue solid) of representative VH domains with VH1-8, VH1-69.2, VH3-48, and VH6-1 germlines (b). Relative thermal stability as measured by DSF of non-stabilized (red bars) or stabilized (blue bars) VHs representing all VH germlines where * indicates an inability to determine the stability due to a lack of expression or clear DSF transition (c).

**Table 1.** Impact of combination designs on the expression and thermal stability of select VH family members.

| VH Germline | VH germline Design | Variant Combination | Fold Change in Expression over wild-type VH | $\Delta T_m$ (°C) |
| --- | --- | --- | --- | --- |
| VH1 (VH1-8) | 1C1 | 17C/82aC 48I | 3.4 | +11.5 |
|  | 1C2 | 17C/82aC 16D 48I | 3.1 | +14.5 |
| VH2 (VH2-5) | 2C1 | 19C/81C 37Y 83T | 1.2 | +16.7 |
|  | 2C2 | 19C/81C 15G 37Y | 1.6 | +11.3 |
| VH3 (VH3-11, 3-20,3-48) | 3C1 | 23C/77C 49A 74E | 2.0, 3.3, 310 | +19.0, +21.5, +17.5 |
|  | 3C2 | 23C/77C 49A 84E | 1.6, 2.0, 69 | +21.0, +22.5, +19.5 |
| VH4 (VH4-39) | 4C1 | 23C/77C 82bD 84P | 11.5 | Nd |
|  | 4C2 | 23C/77C 10T 82bN | 16.1 | +14.2 |
|  | 4C3 | 23C/77C 82bN 84P | 11.5 | +14.3 |
| VH5 (VH5-51) | 5C1 | 28D 48I 83D | 12.3 | $T_m = 76.6^a$ |
| VH6 (VH6-1) | 6C1 | 23C/77C 37Y | 3.5 | +18.3 |
| VH7 (VH7-4) | 7C1 | 17C/82aC | 8.8 | $T_m = 62.5^a$ |

<sup>a</sup>Wild-type VH expression too low to enable measurement

**Table 2.**
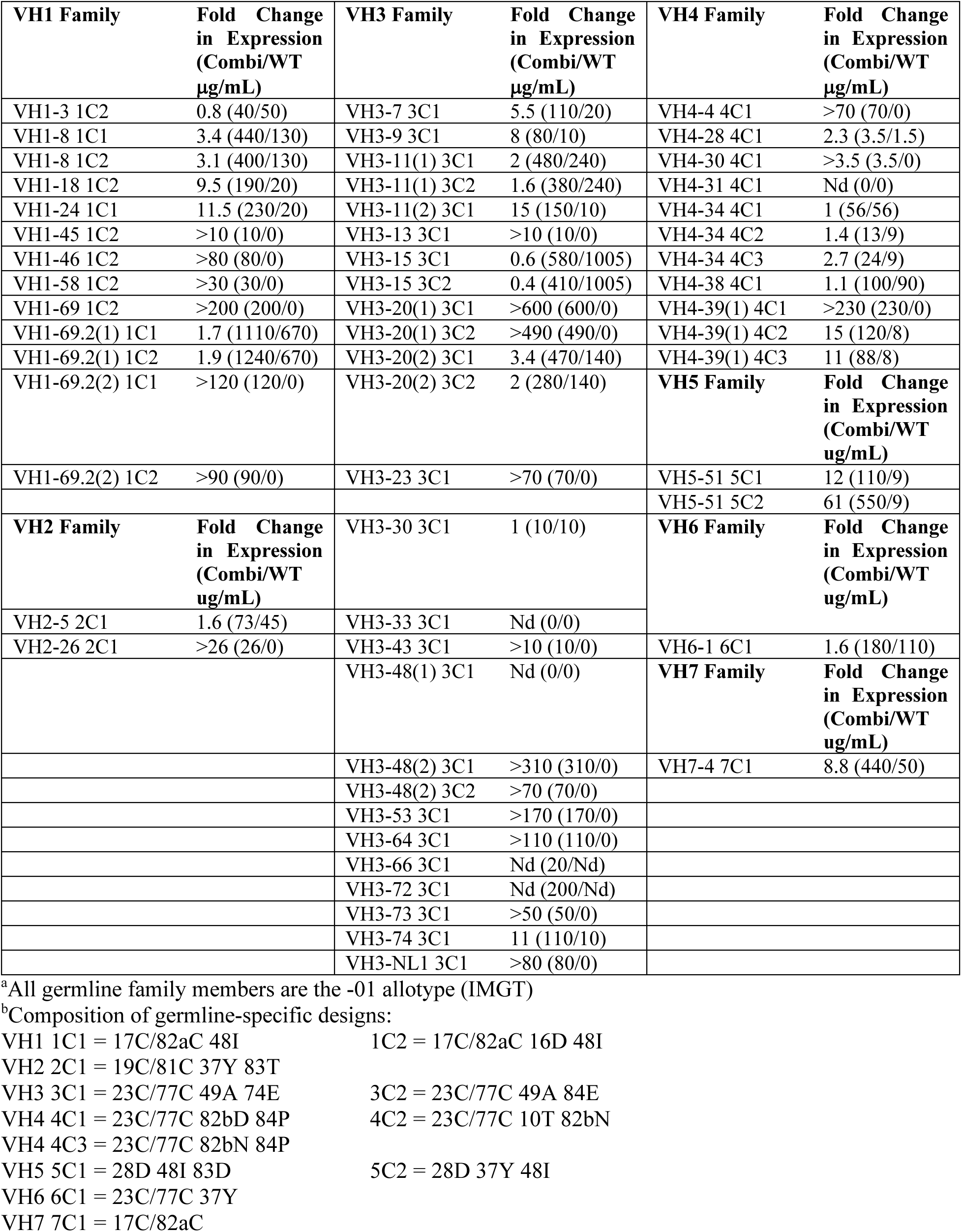
Impact of germline-specific combination designs on the expression of all VH germline family members^a,b^.

### Designs to reduce intrinsic VH/VL and VH/VH interactions

The use of human VH domains as modular subunits requires that intrinsic interactions with antibody light chains (VL domains) be eliminated, along with homotypic VH-VH interactions. Unlike camelid V_HH_ domains whose HCDR3 conformations and framework 2 regions have evolved to maintain a monomeric V_HH_ subunit ^49^, human VH domains have evolved to interact with antibody VL domains. VH affinity for VL depends highly on residues in framework 2 of both the VH and VL, as well as the compositions of HCDR3/LCDR3 ^50^. Binding between VH domains and a control antibody light chain was assessed using flow cytometry by individually expressing various VH domains with N-terminal fusions to human IgG-Fc and C-terminal fusions to the transmembrane (TM) region from PDGFRα on the surface of HEK293 cells in the presence of various secreted light chains. The majority of VH domains did not capture a LC or the capture was very weak; however, a kappa light chain with IGKV1-39/JK4 showed strong binding to a particular VH1-69.2-Fc-TM construct and represented an ideal model system for evaluating the impact of interface modifications on VH/VL affinity. Modifications of VH framework 2 residues 37, 39, 45, and 47, all of which make contacts at the VL interface (**Figure 1a**), were prioritized for screening. Specific modifications 37Y, 39R, and 45E all led to significant reductions in light chain binding without significantly impacting VH-Fc-TM expression (**Figure 3a**). None of these modifications decreased VH stability by DSF and 37Y (as predicted) was typically stabilizing to isolated VH. Other modifications such as 45G, 45A, or modifications of the canonical residue W47 led to a significant loss in VH expression which is likely the result of a decrease in VH domain stability.

**Figure 3.**
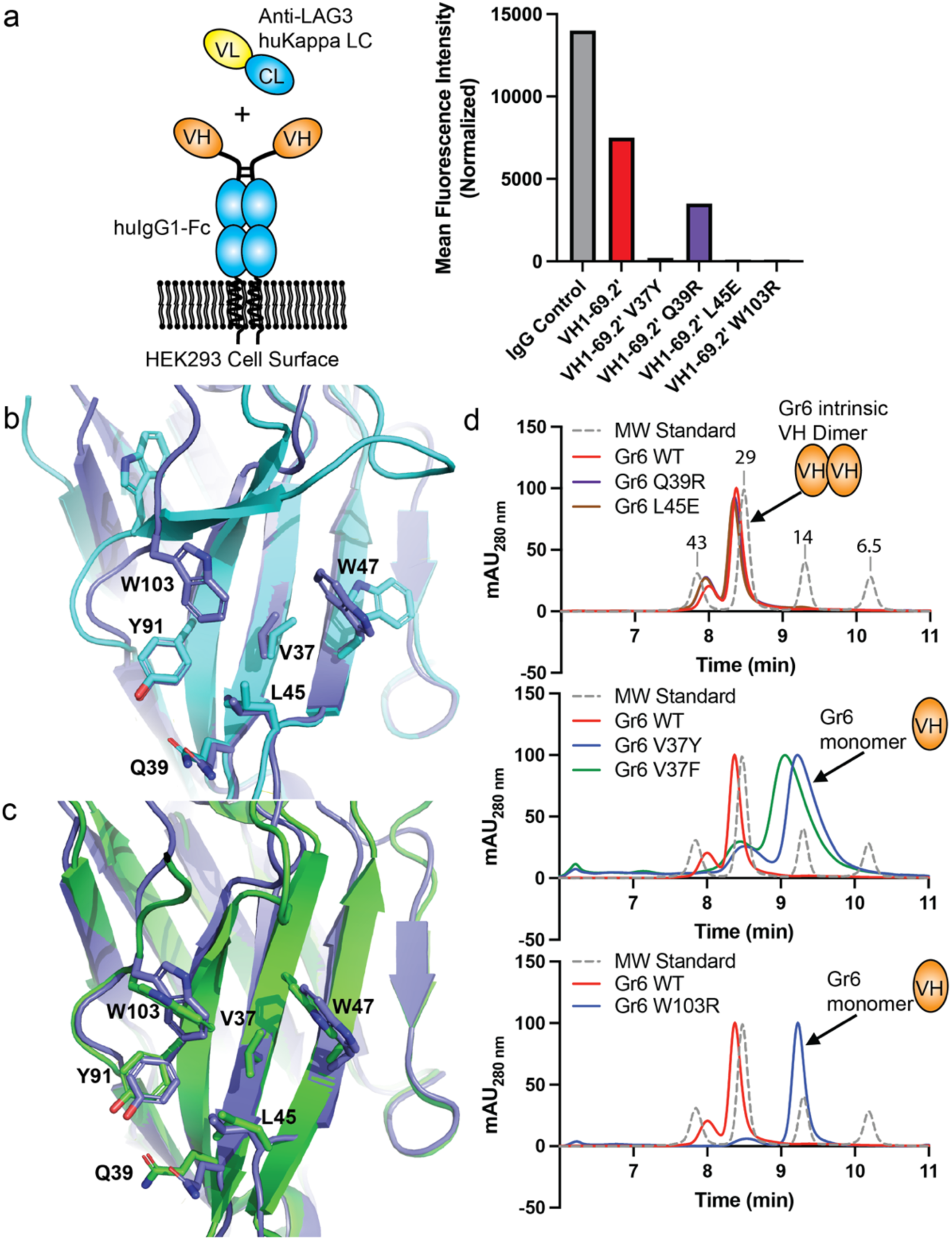
Modification of VL-facing residues to eliminate light chain binding and intrinsic VH homodimerization. Impact of VL-facing amino acid modifications on light chain binding to a control VH1-69.2’ (1C1) VH domain using flow cytometry capture and detection (a). Orientation of VL-facing residues within the VH of a Fab structure (7JOO, cyan) versus that observed within homodimeric Gr6 anti-HER2 (3QYC, purple) (b). Orientation of VL-facing residues within two homodimeric VH structures (3QYC and 1OHQ, purple and green, respectively) suggests a common VH homodimerization motif (c). Impact of VL-facing residue modifications on the ability of the Gr6 VH domain to dimerize according to analytical SEC (d).

Reducing non-native homotypic VH/VH interactions was also considered essential for the use of fully human VHs as combinable modules for creating multi-specific biologics. While VH/VH interactions are less commonly observed than native VH/VL interactions, the high effective concentrations of VHs when combined within a single biotherapeutic poses a risk of VH dimerization and auto-inhibition. Two fully human VH structures deposited in the PDB each form homodimers using the same interface residues as implicated within VH/VL binding (HEL4 binds hen egg white lysozyme and Gr6 binds HER2) ^15,16^. The amino acid conformations within the VH/VH interface are different from what is observed for VH/VL interactions; however, the interface residue conformations within both VH/VH dimer structures are nearly identical, suggesting a conserved motif for VH/VH dimerization (**Figure 3b,c**). The previously described site saturation mutagenesis protocol (Rosetta) was used to identify substitutions at positions 37 and 45 that destabilize the Gr6 VH homodimer but are neutral or stabilizing in the context of a monomer. Modification of valine 37 to phenylalanine or tyrosine was predicted to significantly destabilize the dimer (+9.4 and +11.2 RU, respectively) while stabilizing the monomer (−1.2 and - 1.5 RU, respectively, **Table S4**). Modification L45E was predicted to preferentially destabilize the homodimer as well, but not as significantly (**Table S4**). ThermoMPNN was also used to predict modifications that would impact both VH/VL and VH/VH interactions. Both the V37F/Y and W103R mutations were predicted to reduce the stability of two VH/VL structures and two VH/VH dimeric interfaces, while either stabilizing (V37F/Y) or modestly destabilizing (W103R) the monomeric VH (**Table S4**). The L45E mutation was predicted by both Rosetta and ThermoMPNN to destabilize the dimer preferentially, but with a weaker score than observed for V37F/Y (**Table S4**). Modifications at residues 39 and 44 were not predicted to discriminate significantly between VH/VH dimers and monomers based on ThermoMPNN (**Table S4**).

The Gr6 VH forms a natural dimer and was used as a model system to evaluate different VH/VH disruptor designs via analytical size exclusion chromatography (SEC). The elution times for each variant are provided in **Table S5.** We first evaluated Q39R and L45E as they significantly reduced light chain binding interactions. As predicted computationally by Rosetta and ThermoMPNN, Q39R did not impact VH/VH homodimerization (**Figure 3d**). Modifying G44 to K/Q/R/E also led to no discernable shift towards a monomer peak as predicted by ThermoMPNN (**Table S5**). The V37F and V37Y mutations that were predicted to disfavor homodimerization each led to a near complete shift towards monomer, though there was broadening by analytical SEC suggestive of a residual monomer/dimer equilibrium. Lastly, given that W103 is buried at the center of the interface, we made multiple modifications at this position including W103S/T/R. The W103R modification led to a complete shift from dimer to monomer (**Figure 3d**), while W103S/T had no discernable impact on dimerization. Overall, the V37F/Y and W103R modifications were able to significantly revert the dimer to a monomer. The Gr6 VH stability is very high in part due to stabilization afforded by dimerization and disruption of the dimer tends to be destabilizing. However, addition of 37Y is generally stabilizing to VH domains with weaker dimer propensity as shown for VH6 in **Figure 2b**, which is only stabilized via the 23C-77C disulfide and the 37Y mutation.

### Structural features of the stabilizing designs for each VH germline

To visualize the structural environment of the stabilizing mutations, we solved the crystal structure of the stabilized VH3-20 VH domain and used AlphaFold 2 to generate high-confidence models for design variants from the other germlines (VH1, VH2, VH3, VH4, and VH5)^39^. Attempts were made to crystallize VH1 and VH4 members, but crystals were not observed even after preparing solutions with concentrations above 50 mg/mL. The crystal structure of the anti-IL2Rg VH3-20 domain with the 23C/77C disulfide and S49A and A74E mutations (combination 3C1) was determined at 1.35 Å resolution (**Table S6**). Although the V37Y/W103R modifications developed to occlude homotypic interactions were not incorporated into the stabilized VH3-20 protein at the time of crystallization, the structure was monomeric. The HCDR3 faced the VH/VL and VH/VH interface, which likely occluded homotypic interactions. We also generated a VH3-20 structure using AlphaFold 2. When superimposed, the crystal structure and AlphaFold 2 model were nearly identical with an overall RSMD of 0.37 Å. The tracings of the trailing N-terminus and CDR3 were shifted between the models due to intermolecular crystal packing, though not in a manner leading to any functional consequences. Importantly, AlphaFold 2 predicted the precise orientations of the 23C/77C disulfide and the S49A stabilizing mutation. Given the extremely high fidelity of the AlphaFold 2 model, we used AlphaFold 2 to generate structures for the remainder of the VH germlines to assess the impact of the top designs. A model of the VH1-69.2 domain containing the 3 stabilizing disulfides, each of which were stabilizing alone, illustrates the importance of covalently connecting beta sheets that are distal in primary sequence (B and E strands of the V-class Ig-fold, **Figure 4a**). As shown in **Table S2**, each germline family tended to prefer a specific individual disulfide, though in many cases the domains could be stabilized by alternative single disulfides.

**Figure 4.**
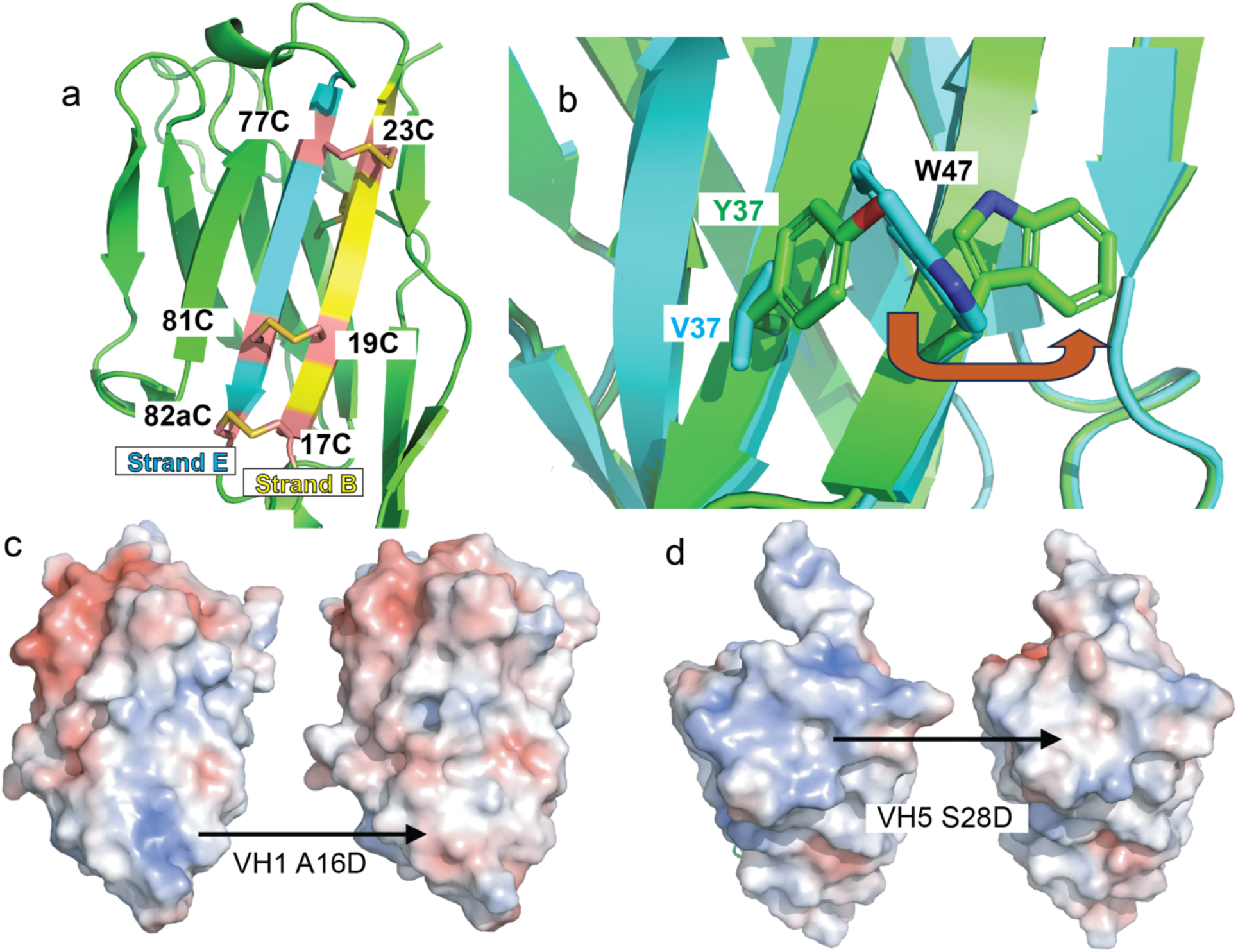
Mechanistic insights into the impact of computational designs on the stability of individual VH domains. AlphaFold2 model of VH1-69.2 depicting the three individually stabilizing disulfides within a single structure to highlight the distance in primary sequence space for the pinned strands (Kabat numbering, only one disulfide inserted per design) (a). Superposition of the crystal structure of one subunit of the Gr6 VH dimer (pdb 3QYC, blue) and the AlphaFold2 model containing the V37Y mutation (green) shows the induced change in the side chain rotamer of W47 that destabilizes both the VH/VL and VH/VH interfaces (b). Electrostatic surfaces depict the neutralization of positively charged patches by the VH1 A16D (c) and VH5 S28D (d). mutations.

Modification of V/I37Y was found to be stabilizing for multiple germline families with the added benefit of reducing both VH/VL heterodimerization and VH/VH homodimerization. AlphaFold 2 was used to evaluate the impact of introducing Tyr at position 37. Mutation to 37Y significantly alters the shape of the former VH/VL and VH/VH interface by pushing 47W into a different rotamer (**Figure 4b**). The VH6-1 and VH7-4 germlines were highly stabilized simply by the addition of the 23C/77C or 17C/82aC disulfides, respectively. VH6-1 stability was also improved by the 37Y modification (VH6 6C1). Additional work with VH7-4 was deprioritized as it is naturally deleted from a large proportion of the human population and could be immunogenic in individuals lacking the gene ^51^.

The impact of the stabilizing designs within the germline families are broken down as follows:

#### VH1 1C2 (17C/82aC A16D M48I)

The A16D mutation places a negative charge between the side chain amines of K10 and K11 (**Figure 4c**), a motif common in all VH1 family germlines as well as in VH5. VH5 naturally contains E16 to neutralize the positive charge cluster. The M84I mutation buries a similar amount of hydrophobic surface area but adopts a backbone orientation well suited to the more constricted β-branched amino acid, reducing the conformational entropy difference between the folded and unfolded states (**Figure S1a**) ^52^.

#### VH2 (19C/81C T15G D83T)

The native T15 adopts a backbone configuration outside the favorable ϕ-backbone dihedral for Thr (+80°). Mutation to Gly should relieve this strain (**Figure S1b**) ^52^. Similarly, D83 adopts a backbone ϕ/ψ configuration abnormal for Asp, but populated by Thr (**Figure S1b**) ^52^. Additionally, mutation to Thr reduces a negative charge patch and the sidechain forms backbone and side chain hydrogen bonds with Asp 86.

#### VH3 3C1 (23C/77C S49A A74E)

The S49A mutation eliminates the burial of a hydroxyl group with unsatisfied hydrogen bonds. Ala at position 49 is the native residue for VH3-30 and VH3-33. The A74E mutation leads to nominal charge balancing commensurate with its nominal impact on expression and stability for two different VH3 family members.

#### VH4 (23C/77C G10T A84P)

Both G10T and A84P likely stabilize the VH domain via reduction of the conformational entropy penalty required to adopt the wild-type conformation as the ϕ/ψ backbone angles of the wild-type residue dihedrals fall within the narrower ranges allowed for the β-branched Thr and Pro (**Figure S1c**).

#### VH5 (S28D M48I)

S28D neutralizes a positively charged patch and forms an additional hydrogen bond with the amide nitrogen of N31 (**Figure 4c**). M48I was also stabilizing for VH5, as it was for VH1, likely due to a reduction in the conformational entropy penalty for forming the wild-type structure (**Figure S1a**).

### Introduction of VH-Select™ into mammalian display libraries leads to enhanced display

Tentarix uses a mammalian display platform, HuTARG™, that exploits natural V-D-J recombination to generate human VH libraries expressed on the surface of HEK293 cells. HuTARG™ libraries of >10^6^ in-frame VH-Fc-TM molecules capture a diverse set of de novo CDR3 sequences in the context of 47 germline/non-stabilized, or 34 VH-Select™ domains that are common throughout the entire human population ^51^. The surface expression levels of non-stabilized human VH domains was compared to two libraries containing VH-Select™ domains incorporating 37Y or 39R/103R as modifications to eliminate intrinsic VH and VL interactions. The non-stabilized VH-Fc-TM library demonstrated a bimodal expression profile with a significant fraction of the library expressing at a much lower level. In contrast, both VH-Select™ libraries comprising 37Y and 39R/103R modifications displayed a more homogenous population at higher levels of VH-Fc-TM expression (**Figure S2**). Profiles of the VH-Select™ domains more closely resembled that of full IgG molecules generated in the HuTARG™ system. We have empirically observed significantly increased numbers of binders emerging from the VH-Select™ libraries.

### The use of VH-Select™ binders for biotherapeutic design

The ultimate goal for stabilizing the human VH germline repertoire is for the discovery of VH-Select™ binders for use as modular binding subunits to be combined within complex multispecific biotherapeutics dubbed ‘Tentacles™’. While antibodies and bispecific antibodies have revolutionized targeted therapy, it remains challenging for these molecules to mitigate off-target toxicities or to deliver multi-pronged activity that may be necessary for optimal efficacy. Tentacles™ bind multiple targets with tunable affinities to achieve the cell and/or tissue specificity required for effective biotherapeutics with enhanced therapeutic index.

Our initial foray into multi-specifics was to generate Tentacles™ by fusing VH binders with low affinity towards broadly-expressed receptors to mAbs with high affinity that drive cell-specificity to achieve conditional activation of a desired subset of immune cells. 41BB and IL2 receptors provide immunological ‘Signals 2 and 3’ necessary for full T cell stimulation upon T cell receptor activation ^53,54^ and have thereby generated significant interest for immunotherapies; however, non-conditional agonism is fraught with systemic toxicity. These side-effects have led to an inability to safely achieve meaningful clinical results for 41BB, ^55^ and limited use for IL2 which was originally approved over three decades ago but is rarely used today ^56^. In an effort to achieve conditional activation of 41BB and IL2 receptors on specific subsets of T cells, we assessed multiple fusion partners including anti-CD8b, anti-PD1 (pembrolizumab/Keytruda®), and anti-LAG3 ^57,58^. CD8b was chosen as a target to specifically expand and activate the cytotoxic T cell compartment to recognize and kill tumor cells. PD1 and LAG3 are known to be upregulated on antigen-specific tumor infiltrating lymphocytes and are good markers for tumor antigen-specific T cells ^59^. Combining low affinity IL2 and 41BB agonists with higher affinity CD8b, PD1, and LAG3 binders should enable cis-conditional 41BB and IL2R T cell activation.

Using VH domains discovered originally as wild-type (non-stabilized) proteins, Tentacles™ with specificity for CD8, IL2Rγ and IL2Rβ were generated with the goal of specifically expanding CD8 effector T cells to bolster natural responses to viruses or tumors without expanding cytotoxic NK or regulatory T cells. After transient expression in HEK293 cells, the behavior of these proteins was clearly non-ideal. Protein A eluates of multiple constructs were typically less than 50% percent pure demonstrating significant aggregation and proteolytic susceptibility that would limit scalability and developability (**Figure 5a**). This result highlights the need for stabilized human VH domains to generate multispecific Tentacles™.

**Figure 5.**
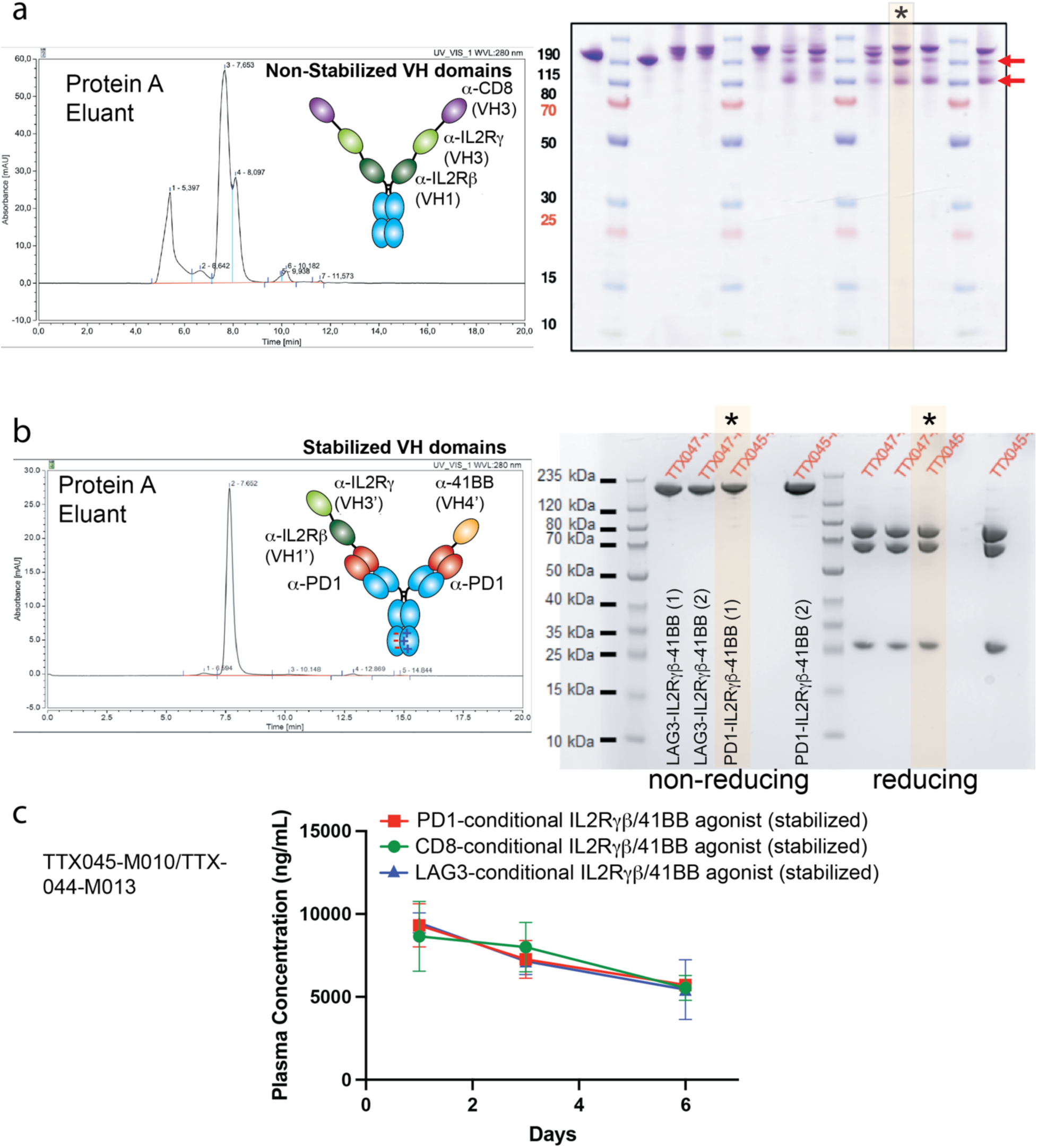
Impact of VH stabilizing designs on the stability and assembly if VH-containing Tentacles^TM^. (a) Analytical SEC and SDS-PAGE characterization of non-stabilized VH-containing multi-specifics post protein A purification. (b) Analytical SEC and SDS-PAGE analysis of VH-containing Tentacles^TM^ that utilize the same IL2Rβ/IL2Rγ VH domains and a 41BB-specific human VH that all contain stabilizing designs. (c) Exposure study of three different Tentacles™ designed for conditional IL2R and 41BB agonism in C57BL/6 mice.

Tentacles™ using VH-Select™ binders were constructed by N-terminal fusion of low affinity anti-41BB and anti-IL2Rβ/IL2Rγc domains to the heavy chains of high affinity anti-CD8b, anti-PD1, and anti-LAG3 mAbs. A modification, N297A, was added to eliminate IgG1-Fc effector function and CH3 heterodimerization designs were added to eliminate excess avidity for the binding of 41BB and IL2Rγ/β ^60^. These optimized Tentacles™ were transiently expressed, purified, and characterized. While not a purely equal comparison with the non-stabilized Tentacles™, these stabilized Tentacles™ contain the same anti-IL2Rγ and anti-IL2Rβ VHs within the non-stabilized Tentacles™ described above. Unlike the non-stabilized Tentacles™, these optimized Tentacles™ showed greater than >85% purity directly following affinity chromatography (**Figure 5b**) and greater than 95% purity after a second column step. The molecules were tested for their ability to demonstrate IgG-like stability *in vivo* by intravenous injection into C57BL/6 mice where no target-mediated drug disposition is expected since none of the targeting arms recognize the mouse receptors. The exposure was tested over six days using the high affinity antigen (CD8, PD1, or LAG3) and anti-human Fc for detection antibody (**Figure 5c**). The exposure was also tested using IL2Rγβ or 41BB antigen to assess the stability of the VH domains, though their lower affinity to these antigens made these ELISAs less sensitive and thus, less accurate. The exposure values using CD8, PD1, or LAG3 antigen, respectively, was generally within 2-fold the values obtained using the VH antigens indicating the intact nature of the Tentacles™ through six days.

Next, the three Tentacles™ (**Figure 6a**) were tested for their conditional agonist activity on T cells. First, the CD8b- and PD1-conditional Tentacles™ were tested for their ability to agonize IL2Rγ/β in a conditional manner on TCR-activated PBMCs. The CD8-conditional Tentacle™ demonstrated dose-dependent activation of CD8+ T cells with a subnanomolar EC_50_, while demonstrating little activity on CD4^+^ T cells or NK cells at concentrations approaching the micromolar range (**Figure 6b**). The PD1-conditional Tentacle™ also demonstrated dose-dependent activation of PD1^+^CD8^+^ and PD1^+^CD4^+^ T cells with subnanomolar EC_50_s, while demonstrating little activity on PD1-NK cells (**Figure 6c**). LAG3 is upregulated on activated CD8, CD4, and NK cells and the LAG3-conditional Tentacle™ activated all three cell types but did not activate T or NK cells that were gated as LAG3 negative (**Figure S3a**). As a control, human IL2 was tested in both experiments and demonstrated a lack of specificity with highly potent activation of all three cell types (**Figure 6a,b**). Conditional 41BB activation was also assessed using the LAG3-conditional Tentacle™ on a 41BB reporter line transfected with full-length LAG3 or PD1 (**Figure 6d and 6e, respectively**). The LAG3-conditional Tentacle™ demonstrated no agonist activity on the PD1-transfected cell line, but strong 41BB agonist activity on the cell line transfected with recombinant LAG3 (**Figure S3a)**. Similar results were observed for the CD8b- and PD1-conditional Tentacles™ (**Figure S3b,c**), though only the LAG3-conditional Tentacle™ contained an affinity matured variant of the 41BB binder with optimal agonist activity. Thus, conditional Tentacles™ demonstrate both IL2R and 41BB agonist activity with high specificity to cells expressing CD8 or PD1.

**Figure 6.**
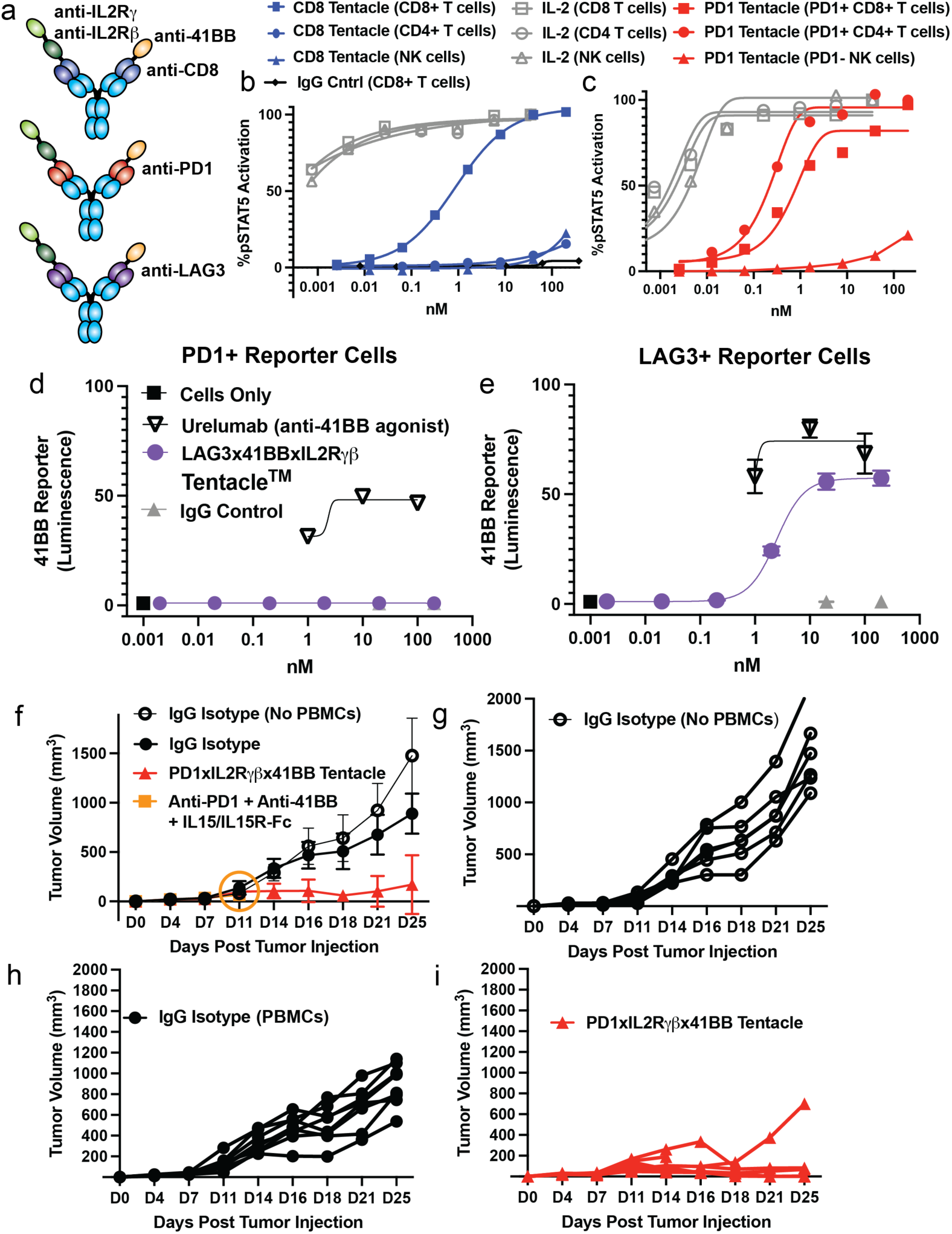
Conditional IL2R and 41BB activity of CD8b- and PD1-conditional agonists. Schematic diagram of the CD8b-, PD1-, and LAG3-conditional IL2R/41BB agonist Tentacles™ (a). Conditional pSTAT5-induction of the CD8b-conditional (left) and PD1-conditional IL2R/41BB Tentacles™ (right) (b). Conditional 41BB agonism of the LAG3xIL2Rx41BB Tentacle on a LAG3-(c) and LAG3+ (d) 41BB-reporter cell line. A375-CMV tumor growth inhibition of the PD1-conditional IL2R/41BB agonist in NSG MHC I/II knock-out mice by the PD1-conditional Tentacle™ (e). A combination of non-conditional PD1 antibody (pembrolizumab), 41BB agonist antibody (urelumab), and IL15/IL15R-Fc was additionally tested, but due to systemic inflammation, the mice lost weight and had to be euthanized per IACUC protocol on day 11 of the study (orange circle). Spider plots tracking tumor growth within the individual animals under each condition are additionally shown (f).

The PD1-conditional Tentacle™ was tested for its anti-tumor activity in a humanized mouse model. HLA-A*02^+^ A375 melanoma cells were transduced to express CMV antigen, which is known to get processed and whose 495-503 peptide (NLVPMVATV) gets bound and displayed by HLA-A*02. Human donor PBMCs were screened for both HLA-A*02 and CMV positivity and a single dual positive donor was selected for engraftment. Other than an IgG control in non-engrafted and PBMC-engrafted mice, a secondary control included the combination of pembrolizumab (PD1 antagonist), urelumab (41BB agonist), and heterodimeric IL15/IL15R-IgG-Fc with ablated Fc-effector function ^61^ to evaluate the impact of a non-conditional trio of biologics with the combined activity of the PD1-IL2Rγβ-41BB Tentacle™.

The PD1-conditional Tentacle™ demonstrated strong anti-tumor activity with improved tolerability (**Figure 6f**). The entire control group treated with non-conditional anti-PD1/anti-41BB/IL15-IL15R-Fc required euthanasia early in the study due to systemic T cell activation and weight loss. The PD1-conditional Tentacle™ showed strong anti-tumor activity with full-tumor regression in 2/6 mice and static tumor growth in all but one mouse. One mouse in the PD1-conditional Tentacle™ group was euthanized due to weight loss. The engrafted PBMC populations were analyzed by flow cytometry and shown to be predominately T cells and to express high levels of PD1 presumably due to residual xeno-responses in the MHCI/II double knock-out strain (**Figure S4**). Thus, even under conditions of unusually high PD1 expression, the PD1-conditional Tentacle™ showed strong anti-tumor activity with improved tolerability over the non-conditional controls.

## Discussion

Here describes the successful design of VH domains across the human genome for improved expression, thermal stability, and monodispersity while abrogating natural interactions with VL and homotypic VH interactions. Improving VH thermal stability has been achieved in specific human VH scaffolds previously. Previous efforts to modify fully human VH domains for use as binder scaffolds have focused on a VH’s ability to refold without aggregating ^14,18,20^ during screening with phage display. These domains generally contained one or more negative charges within HCDR1 that were critical for refoldability. However, refoldability as observed with the anti-IL2Rγ VH binder did not solve all the issues we observed with our multispecifics. Low intrinsic thermal stability led to a susceptibility to proteolysis during lengthy mammalian expression protocols. Disulfides are commonly used to increase the thermal stability of various proteins ^62^. Human VHs have been shown to be stabilized by the addition of disulfides by adding cysteines to positions 49-69 or 35-50 ^24,63^. Non-canonical disulfides also arise naturally in many VHHs including disulfides that stabilize CDR configurations ^64^. We used computational design to identify non-canonical disulfides that broadly stabilize VH structures. We identified several, including the 35-50 disulfide that is commonly observed in rabbit VHs. Disulfides 17-82a, 19-81, and 23-77 stabilized multiple VH domains with specific germline preferences. VH5 was the only germline refractory to all disulfides that were tested. The stabilizing designs for VH5 were entirely based on a combination of point mutations predicted *in silico*. Recently, Kim, Tanha, and coworkers described the use of the 23-77 (24-86 in IMGT numbering) as a means to replace the canonical 22-92 disulfide (23-104 in IMGT numbering) ^63^. Even in the absence of the canonical disulfide, the 23-77 disulfide was able to stabilize a human VH3-11 domain and maintain its resistance to proteolysis (**Figure 5b**). Additionally, computational design was used to specifically find modifications to each germline family that would enable improved, family-specific stability and expression. Given the unique amino acid features of sequences from different germline families, these stabilizing modifications were not generalizable across the different families but were typically applicable within each germline family.

Bispecific and multispecific therapeutic development has advanced considerably over the past decade. These molecules comprised a substantial fraction, roughly one quarter, of antibody-like biologics that entered clinical trials in 2024 ^65–67^. T cell engagers are the most well-known of bispecific antibody approaches that use multiple binders to engage T cells to kill target cells in an antigen-specific context for both cancer and now autoimmunity ^66,68^. Additional approaches to combine inhibitory mechanisms in both autoimmunity and oncology are in clinical development for their ability to combine multiple mechanisms of intervention within a single therapeutic. The utility of camelid VHHs and human VHs to mimic the activity of various cytokines such as IL2, IL12, or 41BBL has been described ^69–72^. However, these modalities require lengthy immunizations and significant engineering to enable ‘clinic ready’ therapeutics. The stabilized fully human VH-domains (VH-Select ™) described here enable rapid and diverse VH discovery using mammalian or phage display without the need for additional protein engineering to make the molecules clinic ready including humanization and deimmunization. The stabilized aspect of VH-Select™ binders also allows for these moieties to be combined and screened *en mass*, without months of stability engineering, to identify unique combinations that achieve novel activity combined with the cellular conditionality required to enable a wider therapeutic index and avoid pleiotropic off-target cell activity. Our initial approach used low affinity human VH-Select™ binders to achieve dual Cis-conditional activation without a substantial increase in the molecular weight of the therapeutic, exemplified by the PD1/LAG3/CD8 x IL2Rγβ x 41BB Tentacles™. Many additional therapeutic approaches can be considered for Tentacles™ that use VH-Select™ binders, which combine an ability to recapitulate the activity of natural growth factors, cytokines, or other signaling proteins with a capacity to manufacture at the scales required for clinical studies. Furthermore, moving outside the cytokine/growth factor space and into other areas that include both conditional agonism and antagonism would unlock additional therapeutic approaches including immune cell redirection or conditional immune checkpoint agonism/antagonism. Overall, we believe the stabilized VH-Select™ binders can be leveraged to solve existing challenges for antibodies and have significant potential to treat human disease.

## Supporting information

Supplement NetMHCPanII4.3 Scoring

## Supplementary Tables and Figures

**Table S1.** High resolution PDB sources for VH domains used for computational analyses.

| Framework | PDB 1 | PDB 2 |
| --- | --- | --- |
| VH1-69.2 | 1RZ7 | 6P9J |
| VH2-5 | 3QRG |  |
| VH3-11 | 6J6Y | 6XKP |
| VH3-15 | 5JR1 | 7JX3 |
| VH3-20 | 1W72 | 7JOO |
| VH3-21 | 6APC |  |
| VH4-39 | 5W6C | 6PZE |
| VH5-51 | 3NAC |  |

**Table S2.** Impact of designed disulfide bonds on the expression and stability of varied germline VHs.

| Disulfide Variant | Expression Ratio (Titer Variant/Titer Wild-Type) |  |  |  |  |  |  |  |  |  |  |  |  |  |  |  |  |
| --- | --- | --- | --- | --- | --- | --- | --- | --- | --- | --- | --- | --- | --- | --- | --- | --- | --- |
|  | VH1-8 | VH1-18 | VH1-69.2 (1) | VH1-69.2 (2) | VH2-5 | VH3-9 | VH3-11 | VH3-15 | VH3-20 (1) | VH3-20 (2) | VH3-30 | VH3-53 | VH4-34 | VH4-39 | VH5-51 | VH6-1<sup>c</sup> | VH7-4 |
| WildType | 1 | 1 | 1 | 1 | 1 | 1 | 1 | 1 | 1 | 1 | 1 | 1 | 1 | 1 | 1 | 1 | 1 |
| 17C 82aC | 2.5 | 1.1 | 1.3 | 2.4 | 0.1 | 1.4 | 1.2 | 1.2 | 0.9 | Nd | 0.70 | 1.1 | 0.1 | 25 | 0.2 | 1.9 | 8.8 |
| 19C 81C | 2.4 | 1.1 | 0.0 | 2.8 | 2.6 | 0.3 | 1.4 | 0.9 | 0.2 | Nd | 0.87 | 0.9 | 0.1 | 16 | 0.1 | 1.6 | 0.4 |
| 23C 77C | 2.3 | 0.2 | 0.9 | 2.1 | 0.3 | 1.1 | 1.1 | 1.1 | 1.1 | 15 | 1.4 | 1.1 | 3.5 | 4 | 0.1 | 2.3 | 1.5 |
| 35C 50C | 2.1 | 0.9 | 1.0 | 1.6 | Nd | 0.9 | 1.1 | 0.6 | 0.9 | 12 | 1.5 | 1.9 | 3.4 | 1 | 0.1 | 2.1 | 10.4 |
| V2C 102C | 0.3 | 0.2 | 0.0 | 3.1 | Nd | 1.9 | 0.6 | 0.4 | 0.5 | Nd | 0.3 | 1.2 | 5.1 | 4 | Nd | Nd | 0.5 |

| Disulfide Variant | Midpoint of Thermal Unfolding (T<sub>m</sub> °C)<sup>a</sup> |  |  |  |  |  |  |  |  |  |  |  |  |  |  |  |  |
| --- | --- | --- | --- | --- | --- | --- | --- | --- | --- | --- | --- | --- | --- | --- | --- | --- | --- |
|  | VH1-8 | VH1-18 | VH1-69.2 (1) | VH1-69.2 (2) | VH2-5 | VH3-9 | VH3-11 | VH3-15 | VH3-20 (1) | VH3-20 (2) | VH3-30 | VH3-53 | VH4-34 | VH4-39 | VH5-51 | VH6-1<sup>c</sup> | VH7-4 |
| WildType | 57 | 65.5 | 62.5 | 64 | Nd | 56 | 73 | 67.5 | 53 | Nd | 57.5 | 72 | Nd | Nd | Nd | 64 | Nd |
| 17C 82aC | 64 | 71.5 | 70 | 71 | Nd | 62 | 74 | 71 | 66.5 | Nd | 72 | 72 | Nd | Nd | Nd | 72 | 62.5 |
| 19C 81C | 62.5 | 69 | Nd | 70.5 | Nd | Nd | 72 | 71 | Nd | Nd | 72 | 72 | Nd | Nd | Nd | 74 | Nd |
| 23C 77C | 61.5 | Nd | 69.5 | 69 | Nd | 64.5 | 72 | 72 | 70 | Nd | 69 | 72 | Nd | Nd | Nd | 77 | Nd |
| 35C 50C | 61 | 70 | 67 | 70 | Nd | 49 | 73 | 64 | 60 | Nd | 72 | 72 | Nd | Nd | Nd | Nd | 67.5 |
| V2C 102C | Nd | Nd | Nd | 71.5 | Nd | 62.5 | 73 | 72.5 | 57 | Nd | 57 | 71 | Nd | Nd | Nd | Nd | Nd |

<sup>a</sup>VHs expressed as VH-Fc. T<sub>m</sub> of IgG1-CH2 was 72 °C, occluding VH T<sub>m</sub> measurements above 72 °C
<sup>b</sup>Nd = Not determined
<sup>c</sup>VH domain contained an 8xhistag instead of huIgG1-Fc

**Table S3.** Impact of single residue modifications on the stability of varied VH germlines.

| Variants | Expression Ratio | Tm<sub>variant</sub> - Tm<sub>WT</sub> (°C) | Variants | Expression Ratio | Tm<sub>variant</sub> - Tm<sub>WT</sub> (°C) |
| --- | --- | --- | --- | --- | --- |
| **VH3-15** | **(Variant/WT<sup>a</sup>)** | **ΔTm** | **VH1-69.2** | **(Variant/WT<sup>d</sup>)** | **ΔTm** |
| A23K | 0.95 | 2.2 | E10Q | 1.3 | 0.6 |
| A23Q | 0.67 | 1.5 | A16D | 0.97 | 3.6 |
| A23Y | 0.74 | 0.8 | A16Q | 1.1 | 0.9 |
| T28D | 0.82 | 3.5 | V37F | 1.2 | 1.6 |
| N31K | 0.88 | 1.0 | M48I | 1.0 | 1.1 |
| A40P | 0.72 | 1.5 | S84E | 0.82 | 1.6 |
| S82bD | 0.74 | 1.2 | S84P | 0.73 | 1.4 |
| T84E | 0.85 | 1.0 | T110V | 0.78 | 4.4 |
| T84P | 0.88 | 0.9 | T110I | 0.71 | 5.6 |
| T110V | 0.60 | 3.8 | **VH4-39** | **(Variant/WT<sup>e</sup>)** | **ΔTm** |
| T110I | 0.62 | 3.8 | Q1E | 1.3 | - |
| **VH3-21** | **(Variant/WT<sup>b</sup>)** | **ΔTm (°C)** | G109Q | 1.3 | - |
| A23Q | 1.3 | - | G10T | 1.5 | - |
| T28D | 3.2 | - | S15G | 1.2 | - |
| T28E | 2.1 | - | S19I | 1.2 | - |
| S33P | 2.0 | - | S54D | 1.1 | - |
| N35A/G/S | 1.4 | - | S82aD | 1.1 | - |
| S49A | 3.0 | - | S82bN | 1.1 | - |
| S52D | 1.8 | - | A84P | 1.1 | - |
| S55E | 1.5 | - | T107I | 1.1 | - |
| Y56E | 2.3 | - | **VH2-5** | **(Variant/WT<sup>f</sup>)** | **ΔTm** |
| A74E | 1.5 | - | T15G | 4.2 | - |
| A84E | 1.5 | - | Q16D | 3.2 | - |
| **VH3-20** | **(Variants/WT<sup>c</sup>)** | **ΔTm (°C)** | I37Y | 3.2 | - |
| V2A | 6 | - | I37H | 2.9 | - |
| A23Q | 3.2 | - | A44D | 4.3 | - |
| T28N | 6.5 | - | A44P | 2.3 | - |
| T28K | 10.6 | - | S65D | 4.9 | - |
| D30K | 7 | - | T73D | 3.4 | - |
| S49A | 145 | - | T73P | 2.5 | - |
| G55E | 11.4 | - | D83L | 2.7 | - |
| A74E | 7.1 | - | D83Q | 2.9 | - |
| N76K | 7.3 | - | D83T | 5.7 | - |
| S77Q | 85.4 | - | P84Y | 4.3 | - |
| A23Q-S77Q | 201 | - | V85R | 3.6 | - |
| **VH5-51** | **(Variants/WT)** | **ΔTm (°C)** | V85R | 3.6 | - |
| S28D | 2.1 | - | V85S | 5.4 | - |
| M48I | 1.7 | - | V85K | 5.0 | - |
| S60D | 1.6 | - | V85T | 6.2 | - |
| S60A | 1.4 | - | T89I | 2.5 | - |
| T68E | 2.1 | - | Q105D | 3.2 | - |
| S76N | 2.2 | - | T107I | 6.1 | - |
| K83D | 4.2 | - |  |  |  |
| A84E | 2.4 | - |  |  |  |

<sup>a</sup>WT = 566 ug/mL<sup>b</sup>WT = 44 ug/mL<sup>c</sup>WT = did not express => arbitrarily assigned 1 ug/mL<sup>d</sup>WT = 187 ug/mL<sup>e</sup>WT = 162 ug/mL<sup>f</sup>WT = 9 ug/mL

**Table S4.** ThermoMPNN ΔΔG° analysis of mutations in VH/VL and VH/VH dimers (PDB IDs: 4LLU/7VYT and 3QYC/1OHQ, respectively)

| Variant | VH/VL $\Delta G^\circ$<br>T-MPNN | VH $\Delta G^\circ$<br>T-MPNN | $\Delta\Delta G^\circ$ | VH/VH $\Delta G^\circ$<br>T-MPNN | VH $\Delta G^\circ$<br>T-MPNN | $\Delta\Delta G^\circ$ | $\Delta\Delta E$<br>Rosetta<br>Units |
| --- | --- | --- | --- | --- | --- | --- | --- |
| W103R | 3.2±0.2 | 2.0±0.2 | 1.2 | 2.5±1.0 | 1.2±1.1 | 1.4 | Nd |
| V37Y | 1.6±0.7 | 0.1±0.6 | 1.5 | 2.7±0.5 | 0.1±0.2 | 2.6 | 11.2 |
| V37F | 1.2±0.4 | -0.1±0.5 | 1.3 | 1.4±0.5 | 0.1±0.02 | 1.3 | 9.4 |
| Q39R | 0.8±0.0 | 0.5±0.1 | 0.3 | 0.7±0.07 | 1.0±0.25 | -0.3 | Nd |
| L45E | 2.2±0.3 | 1.5±0.3 | 0.7 | 2.1±0.6 | 0.6±0.2 | 1.5 | 5.6 |
| W103T | 3.3±0.2 | 2.1±0.2 | 1.2 | 1.7±0.6 | 2.1±0.9 | -0.4 | Nd |
| W103S | 3.2±0.2 | 2.5±0.1 | 0.7 | 1.9±0.3 | 2.1±0.8 | -0.2 | Nd |
| Q105R | 0.5±0.1 | 0.5±0.05 | 0 | 0.7±0.1 | 0.7±0.1 | 0 | Nd |
| Q105K | 0.4±0.1 | 0.3±0.05 | 0.1 | 0.3±0.3 | 0.6±0.3 | -0.3 | Nd |
| Q105E | 0.2±0.2 | 0.1±0.2 | 0.1 | 0.7±0.3 | 0.8±0.5 | -0.1 | Nd |
| G44R | 1.3±0.5 | 1.0±0.3 | 0.3 | 1.7±0.5 | 1.4±0.3 | 0.3 | Nd |
| G44Q | 1.4±0.8 | 0.9±0.2 | 0.5 | 2.0±1.0 | 1.4±0.2 | 0.4 | Nd |
| G44K | 1.4±1.1 | 0.9±0.1 | 0.5 | 2.1±0.7 | 1.6±0.2 | 0.5 | Nd |
| G44E | 1.5±0.7 | 0.9±0.2 | 0.6 | 1.9±0.8 | 1.4±0.1 | 0.5 | Nd |

**Table S5.**
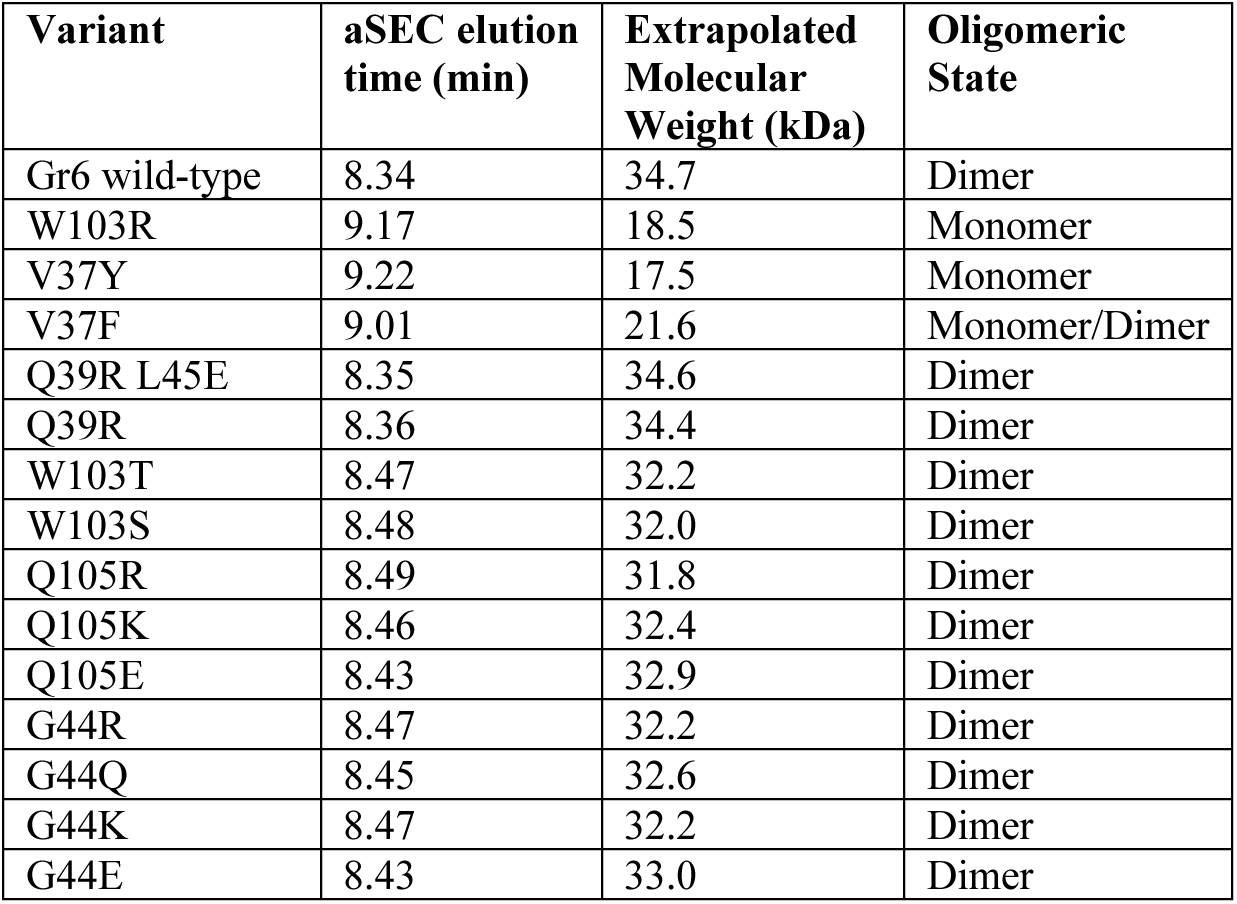
Evaluation of VH/VH interface modifications on dimerization of the anti-HER2 Gr6 intrinsic VH dimer by analytical SEC (aSEC)

| Variant | aSEC elution<br>time (min) | Extrapolated<br>Molecular<br>Weight (kDa) | Oligomeric<br>State |
| --- | --- | --- | --- |
| Gr6 wild-type | 8.34 | 34.7 | Dimer |
| W103R | 9.17 | 18.5 | Monomer |
| V37Y | 9.22 | 17.5 | Monomer |
| V37F | 9.01 | 21.6 | Monomer/Dimer |
| Q39R L45E | 8.35 | 34.6 | Dimer |
| Q39R | 8.36 | 34.4 | Dimer |
| W103T | 8.47 | 32.2 | Dimer |
| W103S | 8.48 | 32.0 | Dimer |
| Q105R | 8.49 | 31.8 | Dimer |
| Q105K | 8.46 | 32.4 | Dimer |
| Q105E | 8.43 | 32.9 | Dimer |
| G44R | 8.47 | 32.2 | Dimer |
| G44Q | 8.45 | 32.6 | Dimer |
| G44K | 8.47 | 32.2 | Dimer |
| G44E | 8.43 | 33.0 | Dimer |

**Table S6.** X-ray crystallography data collection parameters.

|  |  |
| --- | --- |
| **Data reduction** |  |
| Space group | P4<sub>3</sub> |
| *Cell dimensions* |  |
| a, b, c (Å) | 46.821, 46.821, 53.558 |
| $\alpha, \beta, \gamma$ (°) | 90, 90, 90 |
| Resolution (Å)* | 50.0 – 1.35 (1.37 – 1.35) |
| Rmerge* | 0.062 (0.869) |
| Rmeas* | 0.083 (0.784) |
| CC½ (%)* | 0.995 (0.660) |
| I / σI* | 36.5 (1.3) |
| Completeness (%)* | 98.5 (84.6) |
| Total reflections | 154096 |
| Unique reflections* | 25124 (1079) |
| Redundancy* | 6.1 (5.4) |
| Wilson B-factor (Å<sup>2</sup>) | 18.15 |
| VM (Å<sup>3</sup>/Da) | 2.12 |
| **Refinement** |  |
| Resolution (Å) | 33.1-1.35 (1.40-1.35) |
| No. reflections | 24823 (2130) |
| Rwork / Rfree | 18.83 / 20.99 (35.51 / 36.03) |
| *No. atoms* |  |
| Protein | 885 |
| Ligand/ion | 1 |
| Water | 80 |
| *Mean B-factors* |  |
| Overall (Å<sup>2</sup>) | 27.62 |
| Protein (Å<sup>2</sup>) | 27.28 |
| Ion (Å<sup>2</sup>) | 25.38 |
| Water (Å<sup>2</sup>) | 31.36 |
| *R.m.s. deviations* |  |
| Bond lengths (Å) | 0.006 |
| Bond angles (°) | 0.96 |
| *Ramachandran* |  |
| Favored (%) | 99.1 |
| Allowed (%) | 0.9 |
| Outliers (%) | 0 |
| Rotamer outliers (%) | 0 |
| Clashscore | 4.06 |

\* Figures for the highest resolution shell are shown in parentheses.

**Figure S1.**
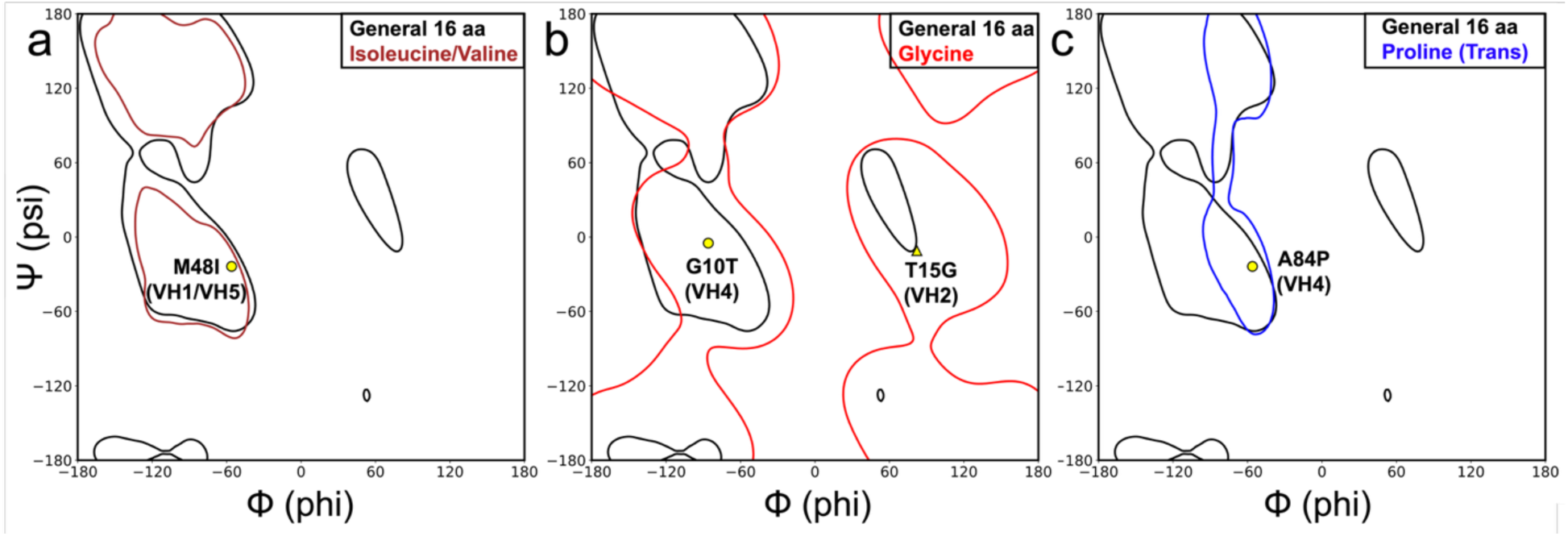
a-c: Ramachandran plots with contours indicating regions that contain 98% of the torsion angles for the specified amino acid(s) from the Top8000 dataset. The Phi/Psi angles of wild type residues for the stabilizing mutations are indicated with yellow markers and derived from PDB ID: 1RZ7, 6PZE, and 3QRG. Mutations M48I, G10T, and A84P (circles) shift to a more restricted conformational landscape and are predicted to reduce the entropy between the folded and unfolded states of their respective VH germlines (a-c). In contrast, the T15G mutation (triangle) relieves torsional strain (b).

**Figure S2.**
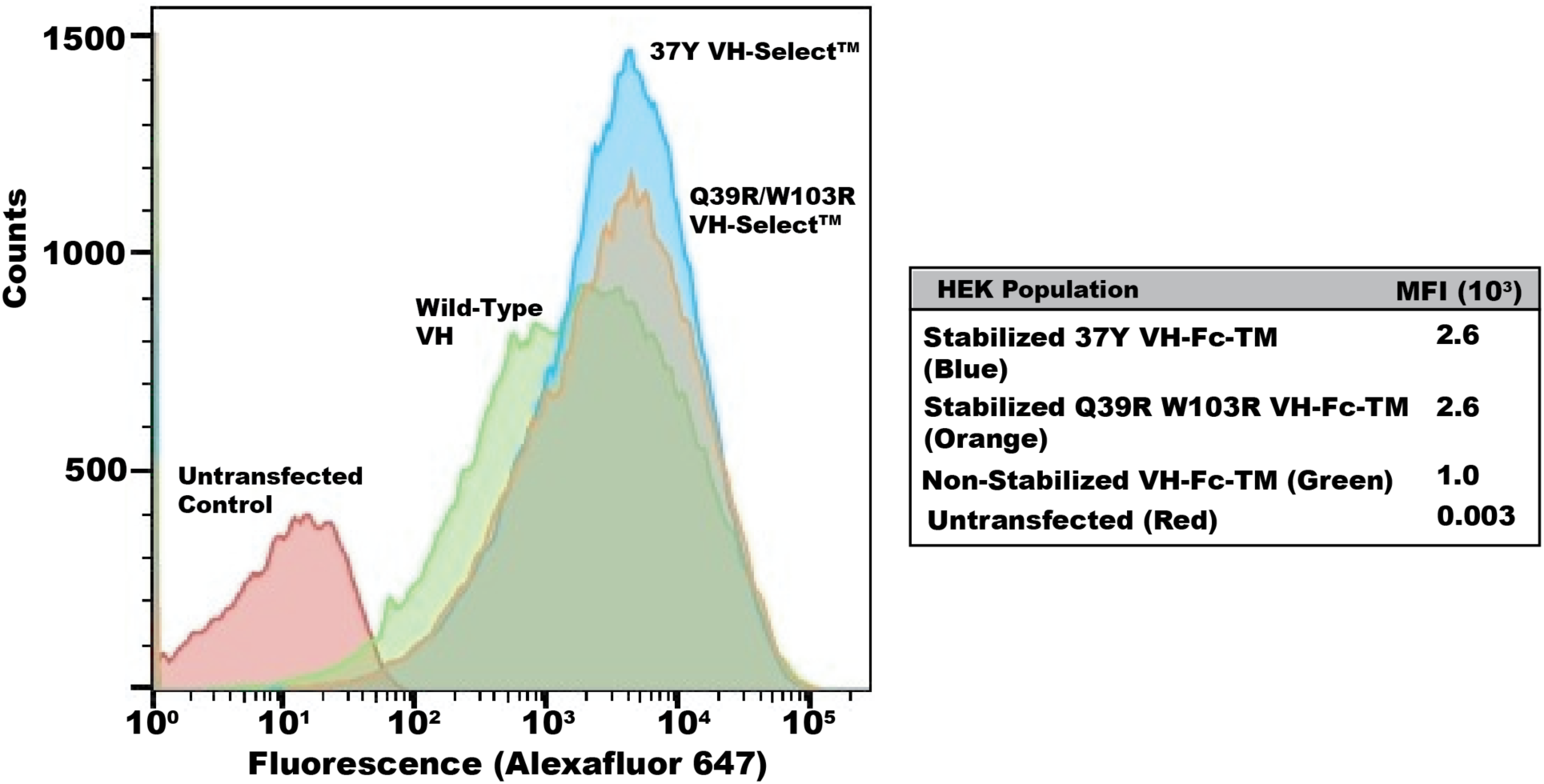
Expression level of recombinant VH-Fc-transmembrane (TM) library with and without stabilizing designs. Fluorescence intensity plots of untransfected HEK293 cells (red), transfected non-stabilized VH-Fc-TM library (green), 37Y VH-Select™-TM library (blue), and Q39R/W103R VH-Select™-TM library (orange). The table on the right provides the mean fluorescence intensity of the populations stained with anti-human-IgG-Fc Alexafluor-647.

**Figure S3.**
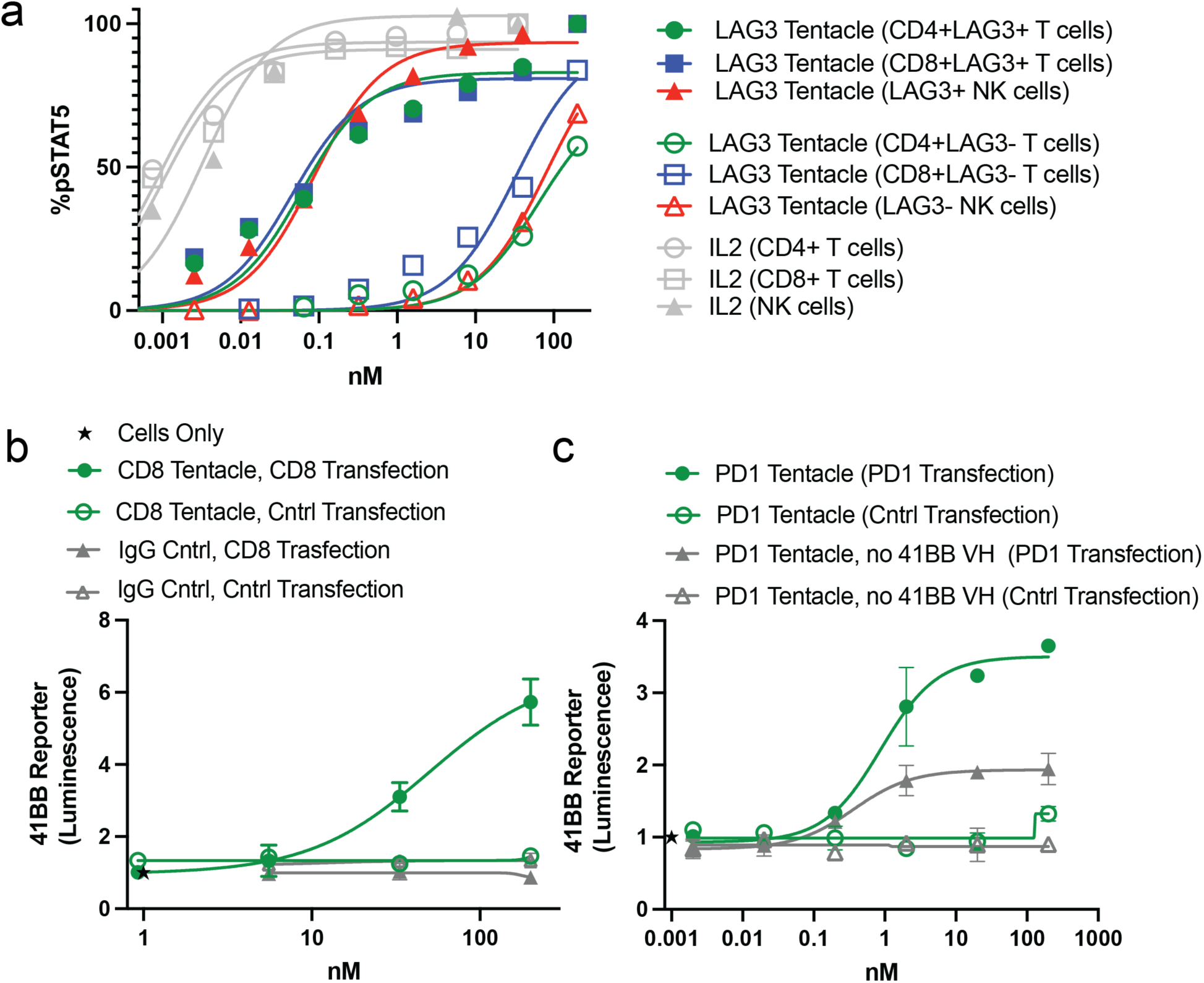
IL2R and 41BB agonism of LAG3 x IL2 x 41BB and CD8b-/PD1-conditional Tentalces, respectively. STAT5 phosphorylation of activated T cells and NK cells by the LAG3 x IL2R x 41BB Tentacle™ (a). Conditional 41BB agonism with the CD8b x IL2R x 41BB (left) and PD1 x IL2R x 41BB (right) Tentacles™ (b).

**Figure S4.**
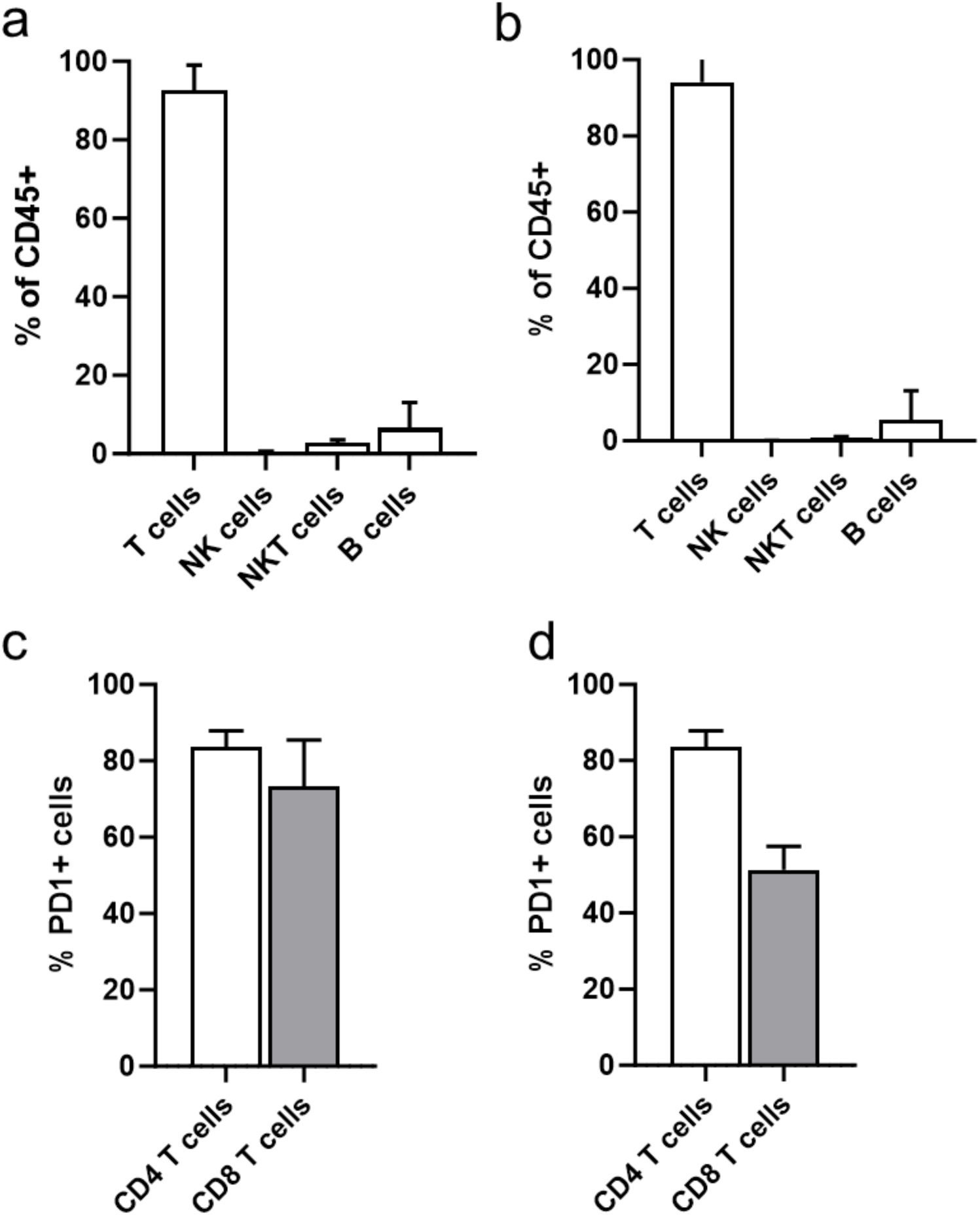
Lineage distribution and PD1 expression of engrafted donor cells in the tumor and blood. Tumors and blood samples were collected on day 25 from the IgG Control group and were stained with various immune markers using flow cytometry to assess the lineage distribution in the tumor (a) and blood (b). PD1 levels in the tumor (c) and blood (d).

