## Supplement NetMHCPanII4.3 Scoring for "Computational Stabilization of the Human VH Germline Repertoire to Enable Conditional Multi-Specific Therapeutic Development"

| VH FAMILY | 17MER (OR MORE) CONTAINING ALL REGISTERS | HLA-DRB1*03:01 HLA-DRB1*07:01 HLA-DRB1*07:01 HLA-DRB1*15:01 HLA-DRB1*15:01 HLA-DRB1*15:01 HLA-DRB5*01:01 | NETMHCPAN 4.3 EL | 9mer | TCR Facing |
|---|---|---|---|---|---|
| **VH1 FAMILY** | | | Score <10 9mer | | |
| | **16D17C** | | | | |
| VH1-2,3,18,24,69,8 | GAEVKKPGDCVKVSCKAS | | 74 | | |
| VH1-45 | GAEVKKTGDCVKVSCKAS | | 39 | | |
| VH1-58 | GPEVKKPGDCVKVSCKAS | | 71 | | |
| VH1-69.2 | GPEVKKPGDCVKISCKVS | | 74 | | |
| | **48I** | | | | |
| VH1-18 | APGQGLEWIGWISAYNG | | 48 | | |
| VH1-2 | APGQGLEWIGWINPNSG | | 22 | | |
| VH1-3 | APGQRLEWIGWINAGNG | | 48I is wild-type | | |
| VH1-8 | ATGQGLEWIGWMNPNSG | | 27 | | |
| VH1-24 | APGKGLEWIGGFDPEDG | | 31 | | |
| VH1-45 | APGQALEWIGWITPFNG | | 48I is wild-type | | |
| VH1-46 | APGQGLEWIGIINPSGG | | 14 | | |
| VH1-58 | ARGQRLEWIGWIVVGSG | | 47 | | |
| VH1-69 | APGQGLEWIGGIIPIFG | | 55 | | |
| VH1-69.2 | APGKGLEWIGLVDPEDG | | 48I is wild-type | | |
| | **82aC** | | | | |
| VH1-2 | ISTAYMELCRLRSDDTA | | 39 | | |
| VH1-3,8,18,24,45,46,58,69,69.2 | ASTAYMELCSLRSEDTA | | 22 | | |
| **VH2 FAMILY** | | | | | |
| | **19C** | | | | |
| VH2-5,70 | LVKPTQTLCLTCTFSGF | | 77 | | |
| VH2-26 | LVKPTETLCLTCTVSGF | | 86 | | |
| | **15G19C** | | | | |

| | | | | | |
|---|---|---|---|---|---|
| VH2-5,26,70 | SGPTLVKPGQTLCLTCTFSGF | | 64 | | |
| | SGPTLVKPGETLCLTCTFSGF | | 63 | | |
| | **81C** | | | | |
| VH2-5,26,70 | TSKNQVVLCMTNMDPVDT | | 24 | | |
| | **81C83T** | | | | |
| VH2-5,26,70 | TSKNQVVLCMTNMTPVDTATYY | | 37 | | |
| **VH3 FAMILY** | | | | | |
| | **23C** | | | | |
| VH3-7,9,11,13,15,20,21,23,30,33,48,53,64,66,73,74,NL | GGSLRLSCCASGFTFSS | | 71 | | |
| VH3-7-> WILDTYPE IS 49A | **49A** | | | | |
| VH3-9, 11 | PGKGLEWVAYISSSGST | HLA-DRB1*15:01 | 6.3 | WVAYISSSG | Partial |
| | | HLA-DRB1*15:01 | 9.9 | WVAYISSSG | Partial |
| VH3-13 | GKGLEWVAAIGTAGDTY | HLA-DRB1*07:01 | 2.3 | LEWVAAIGT | No |
| | | HLA-DRB1*07:01 | 3.6 | LEWVAAIGT | No |
| | | HLA-DRB1*07:01 | 4.3 | LEWVAAIGT | No |
| VH3-20 | PGKGLEWVAGINWNGGS | HLA-DRB3*02:02 | 4.8 | WVAAIGTAG | Partial |
| | | HLA-DRB3*02:02 | 5.3 | WVAAIGTAG | Partial |
| VH3-48 | PGKGLEWVAYISSSSST | HLA-DRB3*02:02 | 6.7 | WVAAIGTAG | Partial |
| | | HLA-DRB1*15:01 | 6.9 | LEWVAAIGT | No |
| | | HLA-DRB5*01:01 | 8.1 | WVAAIGTAG | Partial |
| | | HLA-DRB5*01:01 | 8.3 | WVAAIGTAG | Partial |
| | | HLA-DRB5*01:01 | 8.6 | LEWVAAIGT | No |
| | | HLA-DRB1*15:01 | 9.7 | LEWVAAIGT | No |
| VH3-23 | PGKGLEWVAAISGSGGS | HLA-DRB1*07:01 | 4.4 | LEWVAAISG | No |
| | | HLA-DRB1*07:01 | 5 | LEWVAAISG | No |
| | | HLA-DRB1*07:01 | 5.6 | LEWVAAISG | No |
| | | HLA-DRB5*01:01 | 6.5 | WVAAISGSG | Partial |
| | | HLA-DRB5*01:01 | 8.1 | LEWVAAISG | No |
| | | HLA-DRB1*15:01 | 8.5 | LEWVAAISG | No |
| | | HLA-DRB1*07:01 | 4.4 | LEWVAAISG | No |

| | | | | | |
|---|---|---|---|---|---|
| VH3-30 -> WILDTYPE IS 49A | PGKGLEWVAVISYDGSN | HLA-DRB1*07:01 | 5 | LEWVAAISG | No |
| VH3-33 -> WILDTYPE IS 49A | PGKGLEWVAVIWYDGSN | HLA-DRB1*07:01 | 5.6 | LEWVAAISG | No |
| VH3-43 | PGKGLEWVALISWDGGS | HLA-DRB5*01:01 | 6.5 | WVAAISGSG | Partial |
| | | HLA-DRB5*01:01 | 8.1 | LEWVAAISG | No |
| | | HLA-DRB1*15:01 | 8.5 | LEWVAAISG | No |
| VH3-53 | PGKGLEWVAVIYSGGST | | 19 | | |
| VH3-64 | PGKGLEYVAAISSNGGS | HLA-DRB1*07:01 | 1.7 | LEYVAAISS | No |
| | | HLA-DRB1*07:01 | 1.7 | LEYVAAISS | No |
| | | HLA-DRB1*07:01 | 2.3 | LEYVAAISS | No |
| | | HLA-DRB3*02:02 | 3 | YVAAISSNG | Partial |
| | | HLA-DRB5*01:01 | 3.5 | YVAAISSNG | Partial |
| | | HLA-DRB5*01:01 | 3.9 | LEYVAAISS | No |
| | | HLA-DRB1*15:01 | 4 | LEYVAAISS | No |
| | | HLA-DRB3*02:02 | 4.9 | YVAAISSNG | Partial |
| | | HLA-DRB1*15:01 | 5.1 | LEYVAAISS | No |
| VH3-66 | PGKGLEWVAVIYSGGST | HLA-DRB1*15:01 | 5.3 | LEYVAAISS | No |
| | | HLA-DRB5*01:01 | 5.5 | LEYVAAISS | No |
| | | HLA-DRB3*02:02 | 10 | LEYVAAISS | No |
| VH3-72 | PGKGLEWVARTRNKANS | | 10 | | |
| VH3-73 | SGKGLEWVARIRSKANS | HLA-DRB1*03:01 | 6.4 | LEWVARIRS | No |
| | | HLA-DRB1*03:01 | 6.6 | LEWVARIRS | No |
| | | HLA-DRB1*03:01 | 8.6 | LEWVARIRS | No |
| VH3-74 | PGKGLVWVARINSDGSS | HLA-DRB1*03:01 | 9.2 | LVWVARINS | No |
| | | HLA-DRB1*03:01 | 9.6 | LVWVARINS | No |
| VH3-NL1 | PGKGLEWVAVIYSGGSS | | 18 | | |
| | **74E77C** | | | | |
| VH3-7,9,13,20,21,23,30,33,43,48,53,64,66,NL1 | RFTISRDNEKNCLYLQMNS | VH3-74 | 19 | | |
| VH3-13 | RFTISRENEKNCLYLQMNSL | | 30 | | |

| | | | | | | |
|---|---|---|---|---|---|---|
| | | 74E77C | | | | |
| VH3-72 | RFTISRDDEKNCLYLQMNS | All | | 16 | | |
| VH3-73 | RFTISRDDEKNCAYLQMNSL | | | 12 | | |
| **VH4 FAMILY** | | | | | | |
| | **10T** | | | | | |
| VH4-4,30-1 | VQLQESGPTLVKPSQTL | | | 17 | | |
| VH4-28 | VQLQESGPTLVKPSDTL | | | | | |
| VH4-30-2 | VQLQESGPTLVKPPGTL | | | | | |
| VH4-30-4 | LQLQESGS**T**LVKPSQTL | HLA-DRB1*07:01 | | 6.1 | LQESGSTLV | No |
| | | HLA-DRB1*07:01 | | 9.1 | LQESGSTLV | No |
| VH4-31 | VQLQESGPTLVKPSETL | | | | | |
| VH4-34 | VQLQQWGATLLKPSETL | HLA-DRB1*15:01 | | 2.5 | LQQWGATLL | No |
| | | HLA-DRB1*15:01 | | 3.8 | LQQWGATLL | No |
| | | HLA-DRB1*15:01 | | 5.4 | LQQWGATLL | No |
| VH4-38,59,61 | VQLQESGPTLVKPSETL | | | 17 | | |
| VH4-39 | LQLQESGPTLVKPSETL | | | 22 | | |
| | **23C** | | | 67 | | |
| VH4-4,30-1,30-4,59 | SQTLSLTCCVSGGSISS | | | | | |
| VH4-28 | SDTLSLTCCVSGYSISS | | | 88 | | |
| VH4-30-2 | PGTLSLTCCVSGGSISS | | | 67 | | |
| VH4-31 | SETLSLTCCVSGGSVSS | | | 64 | | |
| VH4-34 | SETLSLTCCVYGGSFSG | | | 90 | | |
| VH4-38 | SETLSLTCCVSGYSISS | | | 88 | | |
| VH4-39,61 | SETLSLTCCVSGGSISS | | | 67 | | |
| | **77C** | | | | | |
| VH4-28 | MSVDTSKNCFSLKLSSV | | >72 | | | |
| VH4-30-2 | ISVDKSKNCFSLKLSSV | | >72 | | | |
| VH4-30-4 | ISVDRSKNCFSLKLSSV | | >72 | | | |
| VH4-4,31,34,38,39,58,61,30-1 | ISVDTSKNCFSLKLSSV | | >72 | | | |
| | **84P** | | | | | |
| VH4-4,28 | SLKLSSVTPVDTAVYYC | | | 20 | | |
| VH4-30-1,30-2,30-4,31,34,38,39,58,61 | SLKLSSVTPADTAVYYC | | | 21 | | |

| | | | | | |
|---|---|---|---|---|---|
| **VH5 FAMILY** | | | | | |
| | **28D** | | | | |
| VH5-51 | ISCKGSGYDFTSYWIGW | | 56 | | |
| VH5-10 | ISCKGSGYDFTSYWISW | | 65 | | |
| | **48I** | | | | |
| VH5-51 | MPGKGLEWIGIIYPGDS | | 43 | | |
| VH5-10 | MPGKGLEWIGRIDPSDY | | 31 | | |
| | **83D** | | | | |
| VH5-51,10 | AYLQWSSLDASDTAMYY | HLA-DRB4*01:01 | 1.6 | LQWSSLDAS | No |
| | | HLA-DRB4*01:01 | 2.5 | LQWSSLDAS | No |
| | | HLA-DRB4*01:01 | 3.6 | LQWSSLDAS | No |
| **VH6 GERMLINE** | **23C** | | | | |
| VH6-1 | SQTLSLTCCISGDSVSS | | 84 | | |
| | **77C** | | | | |
| VH6-1 | INPDTSKNCFSLQLNSV | | 84 | | |
| **VH7 GERMLINE** | **82aC** | | | | |
| VH7-4 | SELKKPGACVKVSCKAS | | 50 | | |
| | **37Y** | | | | |
| VH1-2 | FTGYYMHWYRQAPGQGL | HLA-DRB1*15:01 | 9.3 | MHWYRQAPG | Yes |
| VH1-3,18 | FTSYAMHWYRQAPGQRL | HLA-DRB5*01:01 | 5.3 | WYRQAPGQR | No |
| | | HLA-DRB1*15:01 | 7.5 | MHWYRQAPG | Yes |
| | | HLA-DRB1*07:01 | 7.8 | YRQAPGQRL | Yes |
| | | HLA-DRB1*15:01 | 8.2 | MHWYRQAPG | Yes |
| VH1-8 | FTSYDINWYRQATGQGL | HLA-DRB1*07:01 | 3.6 | YRQATGQGL | Yes |
| | | HLA-DRB1*15:01 | 3.7 | INWYRQATG | Yes |
| | | HLA-DRB1*15:01 | 4.3 | INWYRQATG | Yes |
| | | HLA-DRB1*15:01 | 6 | INWYRQATG | Yes |
| VH1-24 | LTELSMHWYRQAPGKGL | HLA-DRB1*15:01 | 6.7 | MHWYRQAPG | Yes |
| | | HLA-DRB1*15:01 | 7.5 | MHWYRQAPG | Yes |
| | | HLA-DRB1*15:01 | 9 | MHWYRQAPG | Yes |
| VH1-45 | FTYRYLHWYRQAPGQAL | HLA-DRB1*15:01 | 4.8 | LHWYRQAPG | Yes |
| | | HLA-DRB1*15:01 | 5.6 | LHWYRQAPG | Yes |
| | | HLA-DRB1*15:01 | 6.9 | LHWYRQAPG | Yes |

| | | | | | |
|---|---|---|---|---|---|
| VH1-46 | FTSYYMHWYRQAPGQGL | | 11 | | |
| VH1-58 | FTSSAVQWYRQARGQRL | HLA-DRB1*07:01 | 3.1 | YRQARGQRL | Yes |
| | | HLA-DRB1*15:01 | 3.4 | VQWYRQARG | Yes |
| | | HLA-DRB1*15:01 | 4.3 | VQWYRQARG | Yes |
| | | HLA-DRB1*15:01 | 5.8 | VQWYRQARG | Yes |
| VH1-69 | FSSYAISWYRQAPGQGL | HLA-DRB1*15:01 | 2.4 | ISWYRQAPG | Yes |
| | | HLA-DRB1*15:01 | 2.7 | ISWYRQAPG | Yes |
| | | HLA-DRB1*15:01 | 3.8 | ISWYRQAPG | Yes |
| VH1-69.2 | FTDYYMHWYQQAPGKGL | HLA-DRB1*15:01 | 8.6 | MHWYQQAPG | Yes |
| | | HLA-DRB1*15:01 | 9.5 | MHWYQQAPG | Yes |
| VH2-5,26,70 | TSGVGVGWYRQPPGKAL | HLA-DRB1*15:01 | 5.4 | VGWYRQPPG | Yes |
| VH2-5,26,70 | | HLA-DRB1*15:01 | 6.3 | VGWYRQPPG | Yes |
| | | HLA-DRB1*15:01 | 7.9 | VGWYRQPPG | Yes |
| VH3-7 | FSSYWMSWYRQAPGKGL | | 16 | | |
| VH3-9 | FDDYAMHWYRQAPGKGL | HLA-DRB1*15:01 | 6.5 | MHWYRQAPG | Yes |
| | | HLA-DRB1*15:01 | 7.3 | MHWYRQAPG | Yes |
| | | HLA-DRB1*15:01 | 8.8 | MHWYRQAPG | Yes |
| VH3-11 | FSDYYMSWYRQAPGKGL | | 12 | | |
| VH3-13 | FSSYDMHWYRQATGKGL | HLA-DRB1*07:01 | 6.4 | YRQATGKGL | Yes |
| VH3-15 | FSNAWMSWYRQAPGKGL | | 12 | | |
| VH3-20 | FDDYGMSWYRQAPGKGL | HLA-DRB1*15:01 | 8.6 | MSWYRQAPG | Yes |
| | | HLA-DRB1*15:01 | 9.4 | MSWYRQAPG | Yes |
| VH3-21 | FSSYSMNWYRQAPGKGL | HLA-DRB3*02:02 | 8.2 | YSMNWYRQA | Yes |
| VH3-23 | FSSYAMSWYRQAPGKGL | | 11 | | |
| VH3-30 | FSSYAMHWYRQAPGKGL | HLA-DRB1*15:01 | 7 | MHWYRQAPG | Yes |
| | | HLA-DRB1*15:01 | 7.8 | MHWYRQAPG | Yes |
| | | HLA-DRB1*15:01 | 9.3 | MHWYRQAPG | Yes |
| VH3-33,NL1 | FSSYGMHWYRQAPGKGL | | 16 | | |
| VH-43 | FDDYTMHWYRQAPGKGL | HLA-DRB1*15:01 | 6.6 | MHWYRQAPG | Yes |
| | | HLA-DRB1*15:01 | 7.3 | MHWYRQAPG | Yes |
| | | HLA-DRB1*15:01 | 8.9 | MHWYRQAPG | Yes |

| | | | | | | |
|---|---|---|---|---|---|---|
| VH3-48 | FSSYSMNWYRQAPGKGL | HLA-DRB3*02:02 | | 8.2 | YSMNWYRQA | Yes |
| VH3-49 | FGDYAMSWYRQAPGKGL | HLA-DRB1*15:01 | | 9.3 | MSWYRQAPG | Yes |
| VH3-49 | FGDYAMSW**F**RQAPGKGL | | | | | |
| VH3-53,66 | VSSNYMSWYRQAPGKGL | HLA-DRB1*15:01 | | 9.4 | MSWYRQAPG | Yes |
| VH3-64 | FSSYAMHWYRQAPGKGL | HLA-DRB1*15:01 | | 7 | MHWYRQAPG | Yes |
| | | HLA-DRB1*15:01 | | 7.8 | MHWYRQAPG | Yes |
| | | HLA-DRB1*15:01 | | 9.3 | MHWYRQAPG | Yes |
| VH3-72 | FSDHYMDWYRQAPGKGL | | | 14 | | |
| VH3-73 | FSGSAMHWYRQASGKGL | HLA-DRB1*15:01 | | 6.9 | MHWYRQASG | Yes |
| | | HLA-DRB1*15:01 | | 8.1 | MHWYRQASG | Yes |
| | | HLA-DRB1*15:01 | | 9.8 | MHWYRQASG | Yes |
| VH3-74 | FSSYWMHWYRQAPGKGL | | | 11 | | |
| VH4-4 | SGGYYWSWYRQPPGKGL | | >24 | | | |
| VH4-28 | SSSNWWGWYRQPPGKGL | | >24 | | | |
| VH4-30-1 | SGDYYWSWYRQPPGKGL | | >24 | | | |
| VH4-30-2 | SSSNWWSWYRQPPGKGL | | >24 | | | |
| VH4-30-4 | SGGYSWSWYRQPPGKGL | | >24 | | | |
| VH4-31 | SGSYYWSWYRQPPGKGL | | >24 | | | |
| VH4-34,38 | FSGYYWSWYRQPPGKGL | | >24 | | | |
| VH4-39 | SSSYYWGWYRQPPGKGL | | >24 | | | |
| VH4-59 | SGGYYWSWYRQHPGKGL | | >24 | | | |
| VH4-61 | ISSYYWSWYRQPPGKGL | | >24 | | | |
| VH5-51 | FTSYWIGWYRQMPGKGL | HLA-DRB1*15:01 | | 8.6 | IGWYRQMPG | Yes |
| VH5-10 | FTSYWISWYRQMPGKGL | HLA-DRB1*15:01 | | 5 | ISWYRQMPG | Yes |
| | | HLA-DRB1*15:01 | | 6.7 | ISWYRQMPG | Yes |
| | | HLA-DRB1*15:01 | | 8.4 | ISWYRQMPG | Yes |
| VH6-1 | SNSAAWNWYRQSPSRGL | HLA-DRB1*07:01 | | 5.4 | YRQSPSRGL | Yes |
| VH7-4 | FTSYAMNWYRQAPGQGL | HLA-DRB3*02:02 | | 7.4 | YAMNWYRQA | Yes |
| **103R dimer disruptor across all J-chains** | | | | | | |
